# Redefining development in *Streptomyces* bacteria: integrating exploration into the classical sporulating life cycle

**DOI:** 10.1101/2023.08.23.554472

**Authors:** Evan M.F. Shepherdson, Marie A. Elliot

## Abstract

Two growth modes have been described for the filamentous *Streptomyces* bacteria. Their classic developmental life cycle culminates in the formation of dormant spores, where movement to new environments is mediated through spore dispersal. In contrast, exploratory growth proceeds as a rapidly expanding vegetative mycelium that leads to extensive surface colonization and is associated with the release of volatile compounds that promote alkalinization (and reduced iron bioavailability) of its surrounding environment. Here we report that exploratory growth can proceed in tandem with classic sporulating development in response to specific nutritional cues. Sporulating exploration is not accompanied by a rise in environmental pH but has the same iron acquisition requirements as conventional exploration. We found that mutants that were defective in their ability to sporulate were unaffected in exploration, but mutants undergoing precocious sporulation were compromised in their exploratory growth and this appeared to be mediated through premature activation of the developmental regulator WhiI. Cell envelope integrity was also found to be critical for exploration, as mutations in the cell envelope stress-responsive extracytoplasmic function sigma factor SigE led to a failure to explore robustly under all exploration-promoting conditions. Finally, in expanding the known exploration-promoting conditions, we discovered that the model species *Streptomyces lividans* exhibited exploration capabilities, supporting the proposal that exploration is broadly conserved throughout the streptomycetes.

**SIGNIFICANCE:** *Streptomyces* bacteria have evolved diverse developmental and metabolic strategies to thrive in dynamic environmental niches. Here, we report the amalgamation of previously disparate developmental pathways, showing that colony expansion via exploration can proceed in tandem with colony sporulation. This developmental integration extends beyond phenotype to include shared genetic elements, with sporulation-specific repressors being required for successful exploration. Comparing this new exploration mode with previously identified strategies has revealed key differences (e.g. no need for environmental alkalinization), and simultaneously allowed us to define unifying requirements for *Streptomyces* exploration. The ‘reproductive exploration’ phenomenon reported here represents a unique bet-hedging strategy, with the *Streptomyces* colony engaging in an aggressive colonization strategy while transporting a protected genetic repository.

## INTRODUCTION

*Streptomyces* are Gram-positive filamentous bacteria that are ubiquitous in the soil and the sediments of aquatic ecosystems. They are renowned for their specialized metabolism: their products include over two-thirds of antibiotics in clinical use today, alongside a multitude of antifungals, immunosuppressants, pesticides, herbicides, and other bioactive products (1–6). Beyond their impressive biosynthetic capabilities, *Streptomyces* have a complex multicellular, sporulating life cycle. This life cycle begins with spore germination, followed by hyphal tip extension and branching (7, 8). The resulting vegetative mycelium continues growing until reproductive growth initiates, which involves raising unbranched aerial hyphae and subsequently converting these hyphal filaments into chains of dormant spores. Recently, a new growth mode termed ‘exploration’, has been identified as an alternative to the classical sporulating developmental life cycle (9, 10). Exploration is characterized by the rapid outgrowth of vegetative-like hyphae across a solid surface, allowing these canonically non-motile bacteria to colonize areas far from their initial site of inoculation.

Multiple growth conditions are now known to induce exploratory growth, and exploration under these different conditions share several defining characteristics (9, 11). First, exploration is associated with a significantly increased rate of surface area expansion on solid medium, compared with growth during its classical life cycle. Second, exploring colonies develop unique surface morphologies characterized by networks of wrinkles, blisters and buckles that are particularly prevalent near the colony center. And finally, exploring colonies emit trimethylamine, a small basic volatile organic compound that raises the pH of the surrounding area (9, 12). A major consequence of this environmental alkalinization is a corresponding decrease in iron bioavailability (12). This leads to growth inhibition of other microbes in the vicinity and serves as a chemical communication signal that both reinforces the exploration response in the producing *Streptomyces* colonies and promotes the initiation of exploratory behaviour by nearby streptomycetes.

Iron limitation and adaptation to iron scarcity are critical for exploration success. *S. venezuelae* can secrete a suite of desferrioxamine siderophores that assist in surviving their self-imposed iron limiting environment (12). An additional siderophore (foroxymithine) is induced during growth on exploration plates supplemented with glycerol (11). Interestingly, while foroxymithine and desferrioxamine are partially redundant during monoculture growth, the two siderophores exhibit distinct spatial organization, and foroxymithine is critical for effective exploration in competitive culturing conditions (e.g. co-culture with yeast).

Here, we identify a new exploration-promoting condition that induces the growth of *S. venezuelae* in a way that combines characteristics of both exploratory and sporulating growth. This integration of disparate developmental pathways stemmed from the addition of glycerol to classical sporulation growth media. Transcriptional profiling revealed global changes in gene expression between exploration on this glycerol-supplemented medium relative to growth on its unsupplemented, classical development medium. In dissecting the regulation of these sporulating, exploring cells, we observed that the developmental regulators BldD and RsiG were required for wild type growth, as was the extracytoplasmic function (ECF) sigma factor SigE. Unexpectedly, we discovered that exploration under these conditions was not accompanied by a rise in pH. Despite this lack of environmental alkalinization, iron availability and siderophore synthesis remained critical for growth and development, with the alternative foroxymithine siderophore being a key player. Finally, we show that glycerol supplementation of classical growth medium can not only promote the exploration of *S. venezuelae,* but it can also stimulate exploration in the model organism *S. lividans*, for which exploratory growth has not previously been observed.

## RESULTS

### Glycerol supplementation of sporulation-promoting medium stimulates exploratory-like growth

Previous investigations into *Streptomyces* exploration revealed that the addition of glycerol to YP (yeast extract, peptone) medium dramatically enhanced the exploration phenotype (10, 11). Relative to unsupplemented plates, glycerol-grown colonies had an accelerated growth rate, colonized a larger surface area, and developed a more wrinkled surface morphology than previously reported exploring colonies (**Figure 1A**). To gauge whether this glycerol response was confined to cells growing on YP-based media, we inoculated *S. venezuelae* to a growth medium that supports classical sporulating development (mannitol, yeast extract, maltose, or MYM), both with and without added glycerol (MYM and MYMG, respectively). Unexpectedly, growth on MYMG shared characteristics with both classical development and exploration (**Figure 1A**).

**Figure 1.**
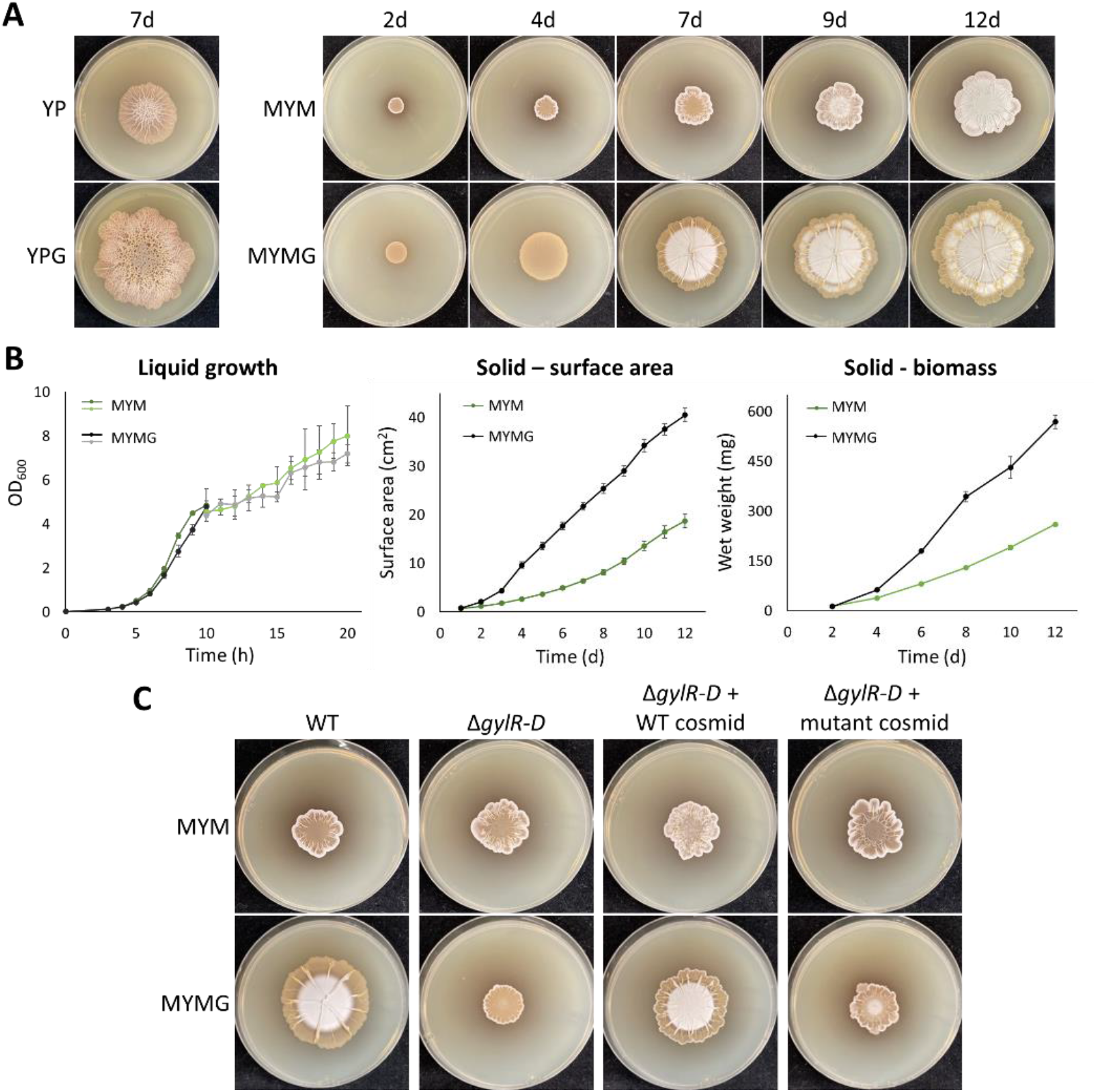
– Growth of *S. venezuelae* on MYMG. **A**) Left: photos of wild type *S. venezuelae* exploring on YP and YPG at 7d of growth. Right: Photos of *S. venezuelae* spotted to MYM and MYMG over the course of 12d of growth. **B)** Quantification of wild type *S. venezuelae* growth rates on MYM and MYMG as measured by OD_600_ in liquid medium (left, three replicates per data point), colony surface area on solid medium (middle, six replicates per data point), and colony wet weight on solid medium (right, three replicates per data point). Error bars represent one standard deviation. Measurements in the liquid growth curve were performed for two sets of flask cultures, one over the first 10h of growth (darker lines) and one over the final 10h of growth (lighter lines). **C)** Wild type *S. venezuelae*, an isogenic glycerol catabolism operon mutant (Δ*gylR-D*), and Δ*gylR-D* complemented with wild type copies of the genes on a cosmid or a vector control (mutant cosmid lacking the *gyl* genes) were spotted to MYM and MYMG and imaged after 7d of growth.

Analogous to supplementing YP with glycerol (YPG), colonies on MYMG expanded faster, yielded larger colonies, and developed a unique wrinkled surface architecture when compared to colonies grown without glycerol (MYM). Intriguingly, these glycerol-grown colonies often formed radially extending wrinkles and displayed concentric rings of aerial hyphae and spores (aerial growth appears as white-ish regions on the plates). Exploration-like colony expansion occurred concurrently with progression through the classical, sporulating life cycle in the central region of the colony (**Supplemental Video 1**). To further investigate the capacity for MYMG to support exploratory-like growth, we generated growth curves for *S. venezuelae* grown on solid or in liquid MYM and MYMG, and compared growth, surface area, and biomass accumulation (**Figure 1B**).

Glycerol supplementation of solid culture stimulated significant increases in surface area and biomass relative to unsupplemented cultures, suggesting enhanced growth rates. These differences were confined to solid culture growth, however, as the differences in growth rates for liquid-grown culture were negligible (**Figure 1B**), suggesting that exploration on solid culture employs a different growth program than in liquid.

To eliminate the possibility that the MYMG phenotype on plates was mediated by altered rheological properties due to glycerol supplementation, we inoculated wild type *S. venezuelae* alongside an isogenic strain in which the operon responsible for glycerol uptake and catabolism was deleted. Growth of the mutant strain on both MYM and MYMG agar resembled that of the wild type on MYM, suggesting that the strain was blind to the presence of glycerol. Importantly, the phenotype could be complemented upon reintroducing the glycerol uptake/catabolic operon (**Figure 1C**). These results suggested that the altered growth pattern of the wild type was due to glycerol metabolism and not to any biophysical properties conferred by glycerol.

We had previously seen that the enhanced exploration phenotype on YPG was specific to glycerol, with no other carbon source having an equivalent effect (11). To determine whether this was also true for MYMG, we tested the behaviour of *S. venezuelae* growing on MYM supplemented with different carbon sources (**Supplemental Figure 1**).

Many of the tested carbon sources (galactose, maltose, mannitol, sorbitol, and sucrose) gave rise to colonies that grew similarly to those without supplementation (MYM), with exceptions being arabinose (similar to glycerol/MYMG), succinate (faster expansion but normal sporulation patterns), glucose (impaired aerial development and exploration), and acetate (no growth).

### Global transcriptional changes are observed between growth on MYM and MYMG

Given the distinct growth characteristics observed for MYMG-relative to MYM-grown cultures, and the similarities of MYMG-grown colonies to exploration, we sought to compare transcription profiles of colonies grown on MYM with or without glycerol supplementation. We isolated and sequenced RNA from colonies grown for 2d (early), 4d (mid) and 7d (late) on MYMG, or 2d and 4d on MYM (Figure 1). Our transcriptomic data revealed global changes in gene expression, both between growth conditions and over time within a single condition. At 4d, 521 genes were differentially expressed (adjusted p-value < 0.05) when comparing MYM– and MYMG-grown cells, while 499 genes were differentially expressed when comparing early and late timepoints for MYMG (**Figure 2A**). Functional categorization of the differentially expressed genes by Clusters of Orthologous Groups (COGs) revealed strong representation among genes associated with primary and secondary metabolism (**Figure 2B**).

**Figure 2.**
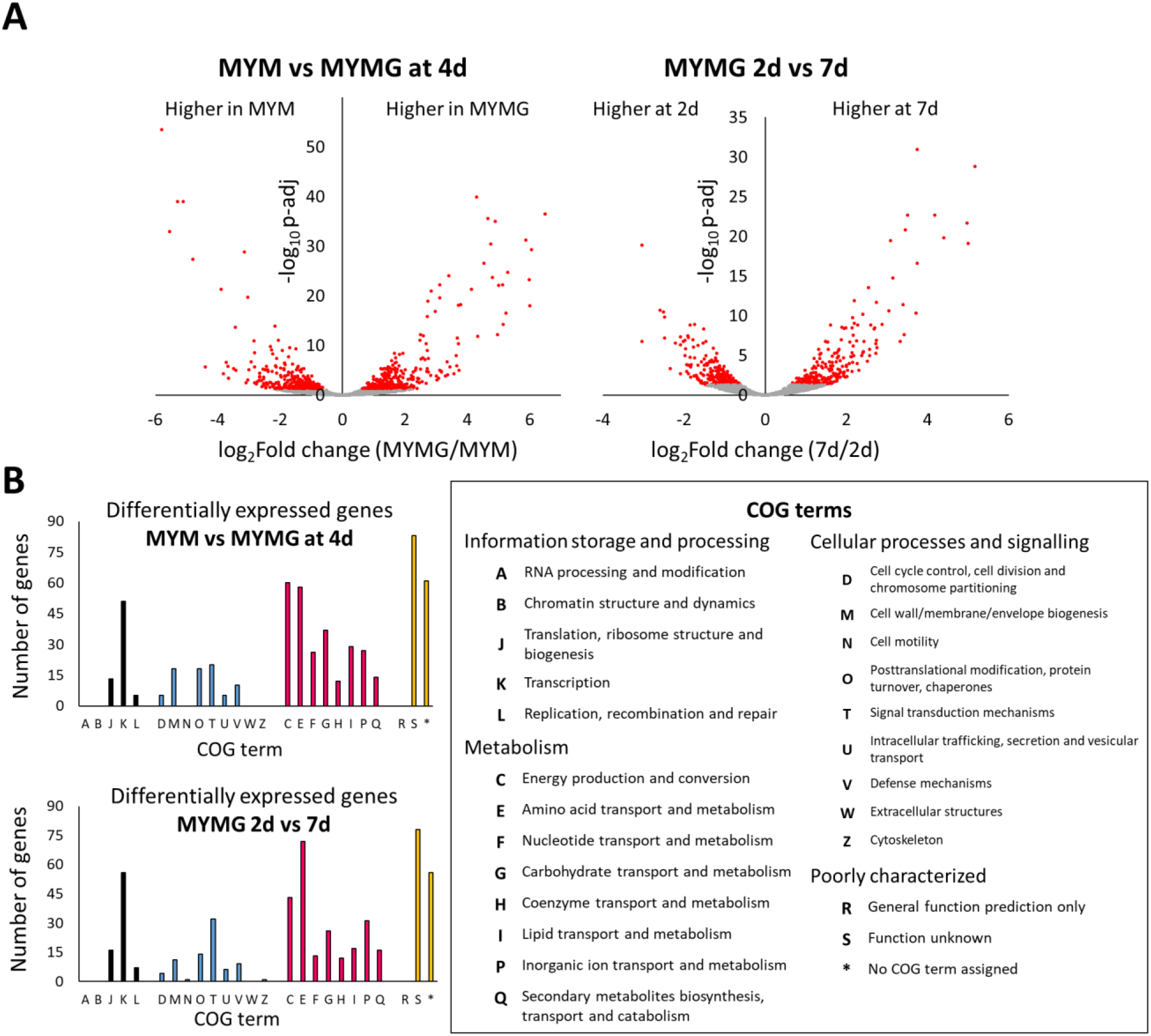
– RNA-sequencing of MYM– and MYMG-grown colonies. **A**) Volcano plots comparing differential gene expression between RNA isolated from MYM and MYMG at 4d of growth (left) and isolated from MYMG at 2d or 7d of growth (right). Data points in red represent genes whose differential expression is statistically significant (adjusted p value < 0.05). **B)** Differentially expressed genes from the conditions identified in **(A)** were sorted according to Clusters of Orthologous Groups (COGs) and plotted according to the number of genes identified within each group.

We found that many of the trends seen when comparing gene expression on YP and YPG were mirrored here (10, 11). We observed changes in expression for genes involved in nitrogen metabolism, central carbon metabolism (glycolysis, TCA cycle, and respiration), regulators of classical development, extracytoplasmic function (ECF) sigma factors, and specialized metabolism/biosynthetic gene clusters (**Supplemental Tables 1** and **2**).

Genes involved in inorganic nitrogen metabolism were among the most significantly differentially expressed genes, and these included genes encoding multiple predicted nitrate/nitrite reductases and transporters. Transcript levels for these genes were higher in MYMG-grown cultures compared to MYM, and their expression increased over time during growth on MYMG. To investigate the possibility that these transcriptional changes reflected an increased demand for nitrogen, we supplemented solid MYM and MYMG with inorganic nitrogen in both fully oxidized (NO_3_) and fully reduced (NH_4_) forms at a range of concentrations (1 µM to 1 mM). Growth was indistinguishable from wild type in all cases, except for the addition of 1 mM nitrate in MYMG which yielded a slightly smaller colony with a bald appearance (lacking aerial hyphae/spores) at 7d of growth (**Supplemental Figure 2**).

A rich source of peptides within the medium has historically been considered a requirement for robust exploration (9). As MYM is not expected to be a peptide-rich medium and since we observed differential expression for genes involved in nitrogen metabolism, we tested the effect of adding more complex nitrogen sources in the form of alanine, casamino acids, and peptone. When added to MYM, we observed a dose-dependent response to all complex nitrogen sources: colonies grew larger, faster, and exhibited surface wrinkling patterns consistent with conventional explorers (**Figure 3**). In contrast, addition of any of these peptide sources to MYMG did not enhance exploration, and instead they inhibited aerial development at all concentrations above 0.1%. This collectively suggested that exploration could be promoted by adding either glycerol or complex nitrogen sources to conventional sporulation-promoting growth medium.

**Figure 3.**
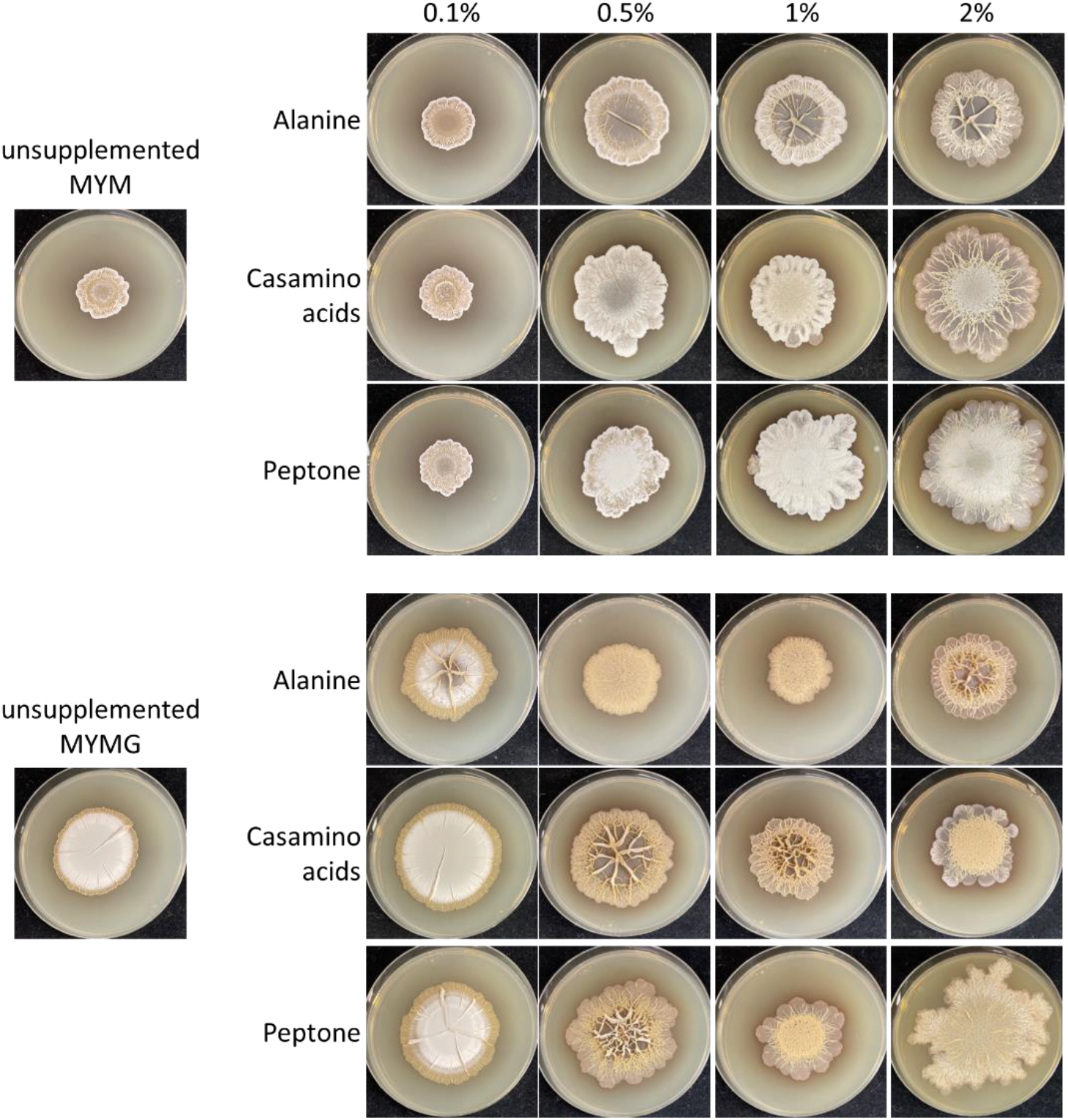
– Effects of complex nitrogen supplementation on MYM and MYMG. Wild type *S. venezuelae* was spotted to solid medium supplemented with different concentrations of alanine, casamino acids, or peptone and photographed after 7d of growth.

### Transcriptional changes in specialized metabolism are not reflected in the bioactivity of MYMG explorers

In addition to observing changes in growth, development and primary metabolism, the transcription of specialized metabolic clusters was also impacted by the addition of glycerol. We found multiple biosynthetic gene clusters were differentially expressed, including those for the antibiotic chloramphenicol and the alternative siderophore foroxymithine, both of which exhibited higher expression during growth on MYMG relative to MYM. We also noted that transcript levels of the venemycin/watasemycin/thiazostatin supercluster increased over time on MYMG. To determine whether these transcriptional differences reflected increased metabolic output in MYMG-grown colonies, we performed antibiotic bioassays using the sensitive indicator bacterium *Micrococcus luteus*. Unexpectedly, MYM-grown cultures yielded more robust inhibition zones compared with MYMG-grown cultures at all timepoints tested (**Supplemental Figure 3**). To determine the nature of the inhibitory molecules, we took advantage of previously constructed strains that were impaired in chloramphenicol and foroxymithine production (11, 13). We found that the majority of antibiotic activity for both MYM– and MYMG-grown cultures was due to the foroxymithine siderophore (**Supplemental Figure 3**).

### MYMG-mediated exploration is associated with a unique response to iron availability

We were surprised to observe high levels of foroxymithine production during growth on MYM and MYMG. We had initially identified foroxymithine as a specialized metabolite that was specifically upregulated during growth on YPG [relative to YP; (11)]. During classical development, we had anticipated that desferrioxamine would be the dominant siderophore due to its broad conservation, and previous characterization in other model streptomycetes (14, 15).

A hallmark of *S. venezuelae* exploration in all previously identified growth conditions was the production of trimethylamine, which raises the pH of the surrounding medium, lowering iron bioavailability. Trimethylamine did not appear to be produced during growth on MYMG (and MYM), as the medium pH remained neutral through 14d of culturing. This raised the question of whether siderophore function was as critical for exploration on MYMG as on other exploration-promoting growth media. We had previously created individual and combined desferrioxamine and foroxymithine mutants (11, 12), and so we used these mutant strains to test the relative importance of these siderophores during growth on both classical growth-promoting MYM and exploration-promoting MYMG. We found that on MYM, growth of the desferrioxamine mutant was indistinguishable from the wild type, while the foroxymithine mutant exhibited reduced aerial development, and the double mutant exhibited a significant growth defect (**Figure 4A** and **Supplemental Figure 4A**). Differences between strains were more pronounced on MYMG: the desferrioxamine mutant displayed wild type growth; the foroxymithine mutant grew slower with reduced aerial development; and the double siderophore mutant did not raise aerial hyphae and failed to expand beyond the original site of inoculation.

**Figure 4.**
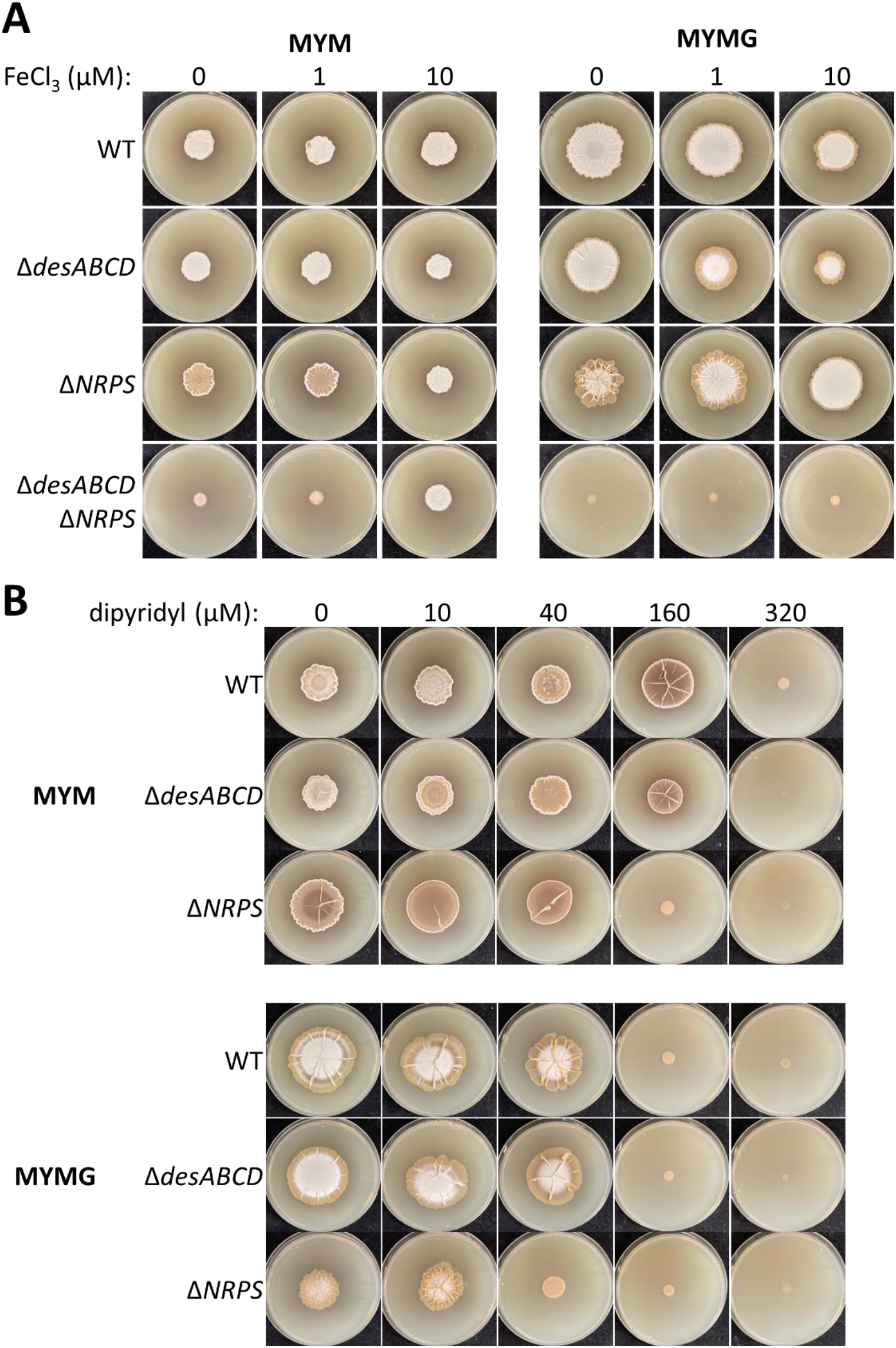
– Effect of iron availability and siderophore repertoire on MYM and MYMG. **A**) A panel of siderophore mutants defective in either desferrioxamine and/or foroxymithine production were spotted to MYM and MYMG supplemented with different concentrations of FeCl_3_. Images were taken after 7d of growth. **B)** Siderophore mutants were spotted to MYM and MYMG supplemented with different concentrations of the iron-specific chelator 2,2’-dipyridyl. Images were taken after 7d of growth.

To evaluate whether the growth deficiencies associated with siderophore loss could be alleviated by iron supplementation, we inoculated the siderophore mutants onto MYM and MYMG supplemented with 1 or 10 µM FeCl_3_. On MYM, 10 µM iron was sufficient to restore wild type growth and development to the foroxymithine and double siderophore mutants (**Figure 4B**). On MYMG, however, increasing concentrations of iron led to decreased colony expansion for both the wild type and desferrioxamine mutant strains, but restored wild type-like growth to the foroxymithine mutant at 10 µM. We found that the double siderophore mutant displayed no growth improvement on MYMG supplemented with 10 µM iron. Consequently, we tested the growth of this mutant using an increased range of iron concentrations (up to 100 µM) and observed that partial phenotypic rescue on MYMG required at least 50 µM iron (**Supplemental Figure 4A**).

We knew from previous work on YPG-mediated exploration that desferrioxamine and foroxymithine exhibited differences in their spatial distribution, with foroxymithine diffusing beyond the colony borders, and desferrioxamine being confined to the colony area (11). We investigated their localization during *S. venezuelae* growth on MYM and MYMG using a Chrome Azurol S (CAS) colourimetric plate assay. CAS overlay plates revealed broad diffusion of siderophore activity in strains producing foroxymithine, whereas activity was limited to colony areas when desferrioxamine was the only siderophore produced (**Supplemental Figure 4B**). Similarly, when excised plugs were tested for activity on CAS agar, large zones of foroxymithine-derived activity were observed for plugs conditioned by wild type *S. venezuelae* on both MYM and MYMG, comparable to zones of activity observed for wild type-conditioned YPG (**Supplemental Figure 4C**). Collectively, these results suggested that increased diffusion of foroxymithine relative to desferrioxamine was a general (pH-independent) property of this siderophore.

As a final test for how iron dynamics affected growth on MYM and MYMG, we lowered the concentration of bioavailable iron by adding increasing concentrations of the iron-specific chelator 2,2’-dipyridyl (**Figure 4C**). Growth on MYMG was more sensitive to 2,2’-dipyridyl, with wild type growth being impacted at 40 µM, while on MYM, 160 µM was required to affect growth, suggesting a greater demand for iron under exploring conditions. For the siderophore mutants, the desferrioxamine mutant resembled the wild type at all concentrations tested, while the foroxymithine mutant was at least 4-fold more sensitive than the wild type to iron limitation on both MYM and MYMG, reinforcing the importance of this siderophore to both classical development and exploration.

### Regulators of classical development contribute to exploration on MYMG

Given the interesting sporulation characteristics of the wild type strain during exploration on MYMG, we wanted to determine whether development and exploration were co-regulated under these growth conditions, as no strong connections had been observed previously (9, 16). We inoculated a collection of classic developmental mutants (predominantly *bld* and *whi* mutant strains that are defective in aerial hyphae formation or sporulation, respectively; a schematic of their relative positions within the regulatory hierarchy is provided in **Figure 5A**) to a panel of media including MYM, MYMG, YP, and YPG, to assess whether these gene deletions perturbed growth under distinct sporulation– and exploration-promoting conditions (**Figure 5B**).

**Figure 5.**
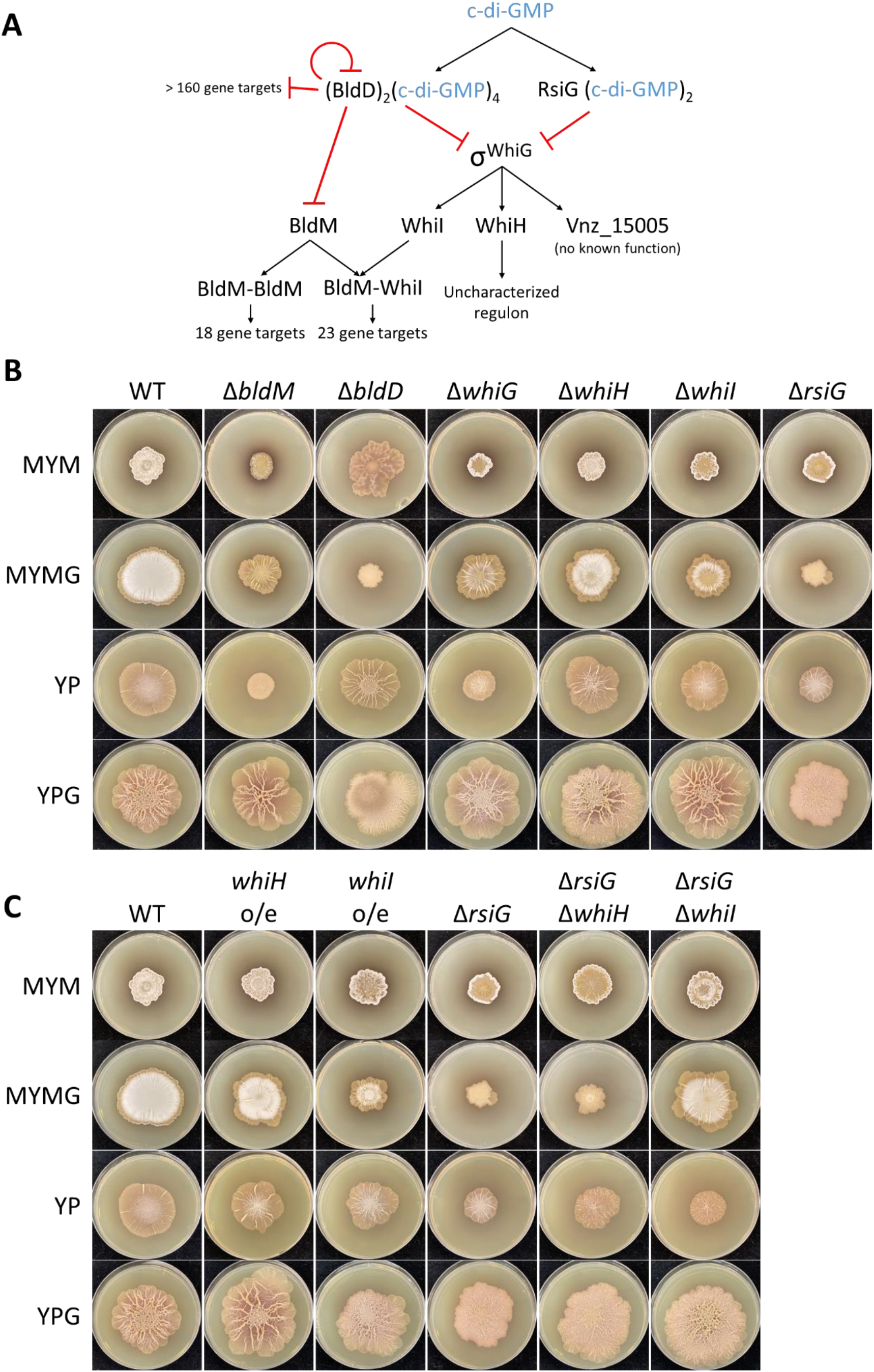
– Regulators of classical development affect exploration on MYMG. **A**) Deletion strains for select regulators of sporulating development were spotted to MYM, MYMG, YP, and YPG and photographed after 7d of growth. **B)** The contribution of *whiH* and *whiI* to normal development across different media types was further examined by spotting strains expressing each gene from a strong constitutive promoter (*whiH* o/e and *whiI* o/e) and by deleting each gene within a Δ*rsiG* mutant background. Images were taken after 7d of growth.

Most developmental mutants exhibited exploratory growth on YP and YPG, with *bldM, whiG* and *rsiG* showing expansion delays on YP relative to wild type, and all showing some differences in surface architecture. In contrast, on MYMG, their growth partitioned into two categories. Mutations that blocked development (Δ*bldM*, Δ*whiG*, Δ*whiH*, and Δ*whiI*) had little impact on exploratory growth on MYMG, with colonies also developing wrinkling patterns like that of the wild type. However, mutations that accelerated development and led to hypersporulation (Δ*bldD* and Δ*rsiG*) abrogated exploration on MYMG.

To gain insight into the mechanism underlying this exploration repression, we focused our attention initially on RsiG, as its downstream regulatory network is much less complex than that of BldD (**Figure 5A**). RsiG is the cognate anti-sigma factor of the sporulation sigma factor WhiG (17, 18). WhiG directs the transcription of three genes in *S. venezuelae*: the developmental regulator-encoding genes *whiH* and *whiI*, and a poorly characterized hypothetical protein-encoding gene *vnz_15005* (17). Thus, in a *rsiG* mutant, we expect precocious WhiG activity and premature/hyper-expression of its target genes.

To determine whether a WhiG target gene(s) was responsible for the *rsiG* phenotype, we independently cloned *whiH* and *whiI* downstream of a strong constitutive promoter (*ermE\**). The resulting constructs were introduced into a wild type background to test whether either could recapitulate the Δ*rsiG* phenotype (**Figure 5B**). In parallel, we created *whiH* and *whiI* deletions in the Δ*rsiG* background to test whether eliminating the downstream targets of RsiG could rescue exploratory growth on MYMG (**Figure 5B**).

We found that constitutive, high-level *whiH* expression in a wild type background had little effect on the exploration phenotype on any medium type, while equivalent expression of *whiI* in the same background impaired growth on MYMG. Consistent with these observations, deleting *whiH* in the Δ*rsiG* background had no impact on the *rsiG* mutant phenotype on MYMG (and other tested media), whereas a Δ*rsiG* Δ*whiI* double mutation rescued the *rsiG* exploration defect (**Figure 5B**). Together, these data strongly implicate early overexpression of *whiI* as being detrimental to exploratory growth on MYMG.

WhiI associates with BldM to form a BldM-WhiI heterodimer that activates the expression of late-sporulation specific genes (19) (BldM can also form a homodimer that controls an independent set of genes; WhiI does not appear to function on its own). Given this, we wondered whether overexpressing *bldM* would have the same effect as overexpressing *whiI*. We found that unlike the situation with *whiI,* overexpressing *bldM* had no significant effect on exploration (**Supplemental Figure 5**). This suggested that either WhiI may act independently of BldM during exploration, or that premature assembly of the BldM-WhiI heterodimer adversely affects exploration. We took advantage of previous chromatin immunoprecipitation (ChIP) sequencing analyses of BldM and WhiI, and identified one target gene (*murA2*) that appeared to be bound more strongly by WhiI than BldM based on ChIP enrichment scores, making it a plausible candidate for WhiI-specific control (19). However, we found that overexpressing *murA2* in a wild type background did not recapitulate the *whiI* overexpression phenotype (**Supplemental Figure 5**). This suggested that it was not responsible for the effect observed, and that the exploration defect may be the result of premature BldM-WhiI assembly, although we cannot formally exclude the possibility that during exploration, WhiI acts alone to control an independent set of genes.

### The cell-envelope responsive sigma factor SigE is required for exploration under diverse growth conditions

In addition to a role for developmental regulators in controlling sporulation-associated exploration, we also identified multiple ECF sigma factors whose genes were differentially expressed (both over time for MYMG-grown cultures, and between MYM and MYMG cultures), including the broadly conserved *sigQ* (**Supplemental Tables 1 and 2**). *sigQ* had previously been identified as a differentially expressed gene in earlier exploration (YP/YPG) RNA-seq datasets (10). We tested the exploration capabilities of *sigQ* mutant and overexpression strains on MYMG, and observed only subtle differences relative to wild type during growth on either MYM or MYMG (**Supplemental Figure 6**). The ECF sigma factor *sigE* had also emerged as a target of interest in expression analyses of YP– and YPG-grown exploring cultures. In *Streptomyces*, SigE is responsible for maintaining cell envelope integrity, and its activation upon sensing envelope stress by the two component system CseBC leads to the upregulation of a large regulon in *S. coelicolor*, many of which encode cell wall– or cell envelope-associated enzymes (20–23). As the rapid growth rate associated with exploration may require greater investment to maintain cell envelope integrity relative to slower growing classically developing colonies, we tested the effects of overexpressing and deleting *sigE* under a range of exploration growth conditions. In contrast to *sigQ*, multiple independently-generated *sigE* deletion mutant strains were found to be significantly impaired in exploration on YP, YPG and MYMG (**Figure 6**), but grew well on the classical development medium MYM. Overexpressing *sigE* had minimal effects. Curiously, introduction of the overexpression construct into any of the Δ*sigE* mutants could fully complement growth of the mutant on MYMG but not on YP or YPG.

**Figure 6.**
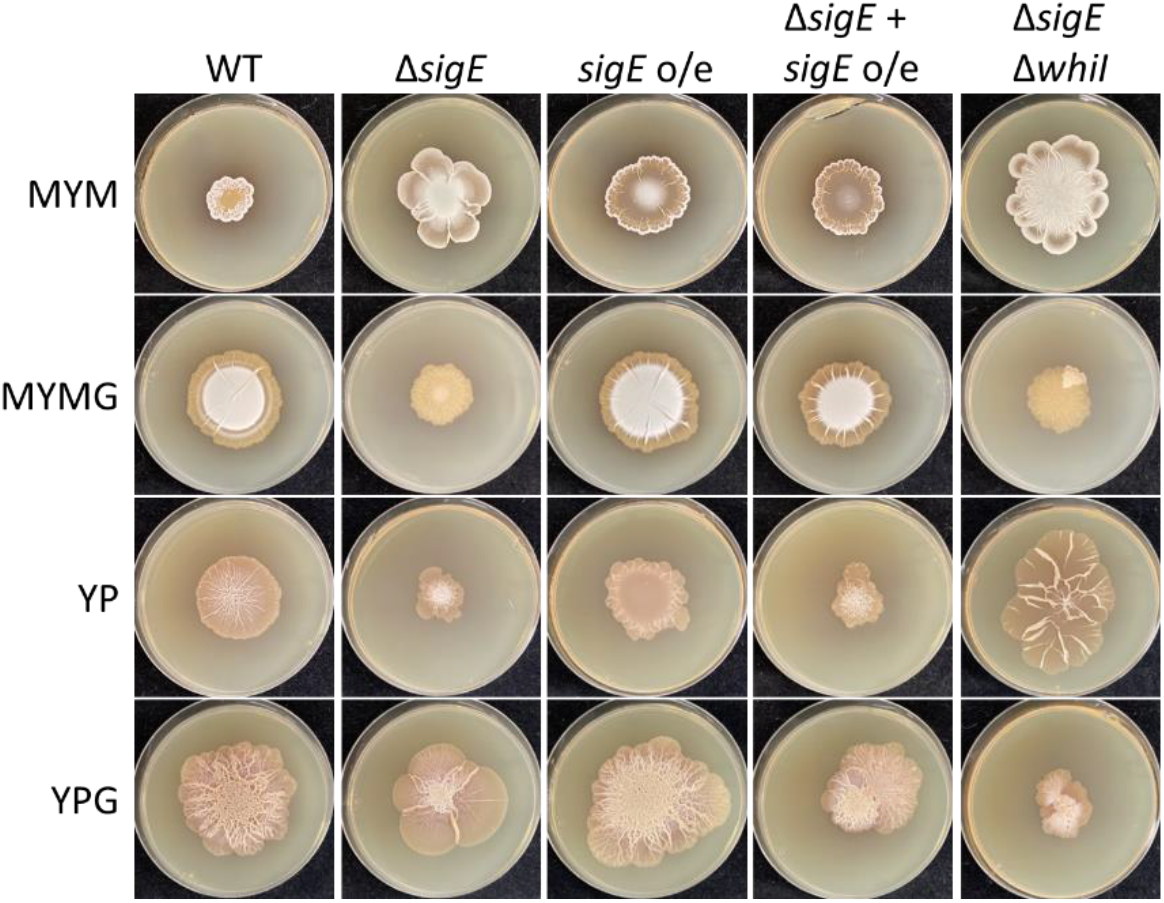
– SigE function is important for robust exploration. Strains of *S. venezuelae* in which the cell envelope stress-responsive extracytoplasmic sigma factor *sigE* was deleted or placed under a strong constitutive promoter (*sigE* o/e) were spotted to different media. A complemented strain (where *sigE* was cloned under the control of the strong constitutive *ermE\** promoter and introduced into the deletion mutant; Δ*sigE* + *sigE* o/e) and a combined double deletion mutant for *sigE* and *whiI* were also spotted to the same media. Images were taken after 7d of growth.

On YP and YPG, but not MYMG, the Δ*sigE* mutant consistently acquired suppressor mutations that resulted in irregular, rapidly exploring outgrowths extending from the main body of the colony. As our investigation into the effects of classical developmental regulators revealed that inappropriate signalling downstream of WhiI negatively impacted exploration on MYMG, we tested whether deleting *whiI* in the Δ*sigE* background could rescue the MYMG phenotype as it had for Δ*rsiG*. We found that while the Δ*sigE* Δ*whiI* mutant was effectively indistinguishable from the *sigE* mutant when grown on MYMG and YPG, exploration was rescued – and even enhanced albeit with very different colony architecture – during growth on YP (**Figure 6**; final column). These observations collectively suggested that exploration is profoundly impacted by SigE, and that there may be distinct downstream regulatory programs that are activated depending on the growth conditions.

### MYMG can promote exploration in species that do not explore under conventional exploration conditions

When exploration was first described, it was reported that ∼10% of wild *Streptomyces* isolates (out of 200) could explore when grown in co-culture with yeast (9). A key question has been whether exploration capabilities are confined to a subset of streptomycetes, or if it is widespread but the initiating stimuli had yet to be identified. We set out to investigate whether our increasingly diverse panel of exploration-promoting growth conditions could stimulate exploratory growth in model *Streptomyces* species not known to explore. We focused our attention on *S. coelicolor* and *S. lividans*, and inoculated them on MYM, MYMG, YP, and YPG (**Figure 7A**). Consistent with previous studies, after 14d of growth, no exploration was observed for either strain on YP or YPG, and robust sporulation was observed for both on MYM. On MYMG, however, *S. lividans* appeared to be exploring: the colony expanded to cover a much larger area of the plate than when grown on MYM, and it developed a complex network of wrinkles with a highly structured core (**Supplementary video 2**). Screening of wild *Streptomyces* from the Wright Actinomycete Collection revealed that additional isolates exhibited exploratory-like growth on MYMG (**Figure 7B**).

**Figure 7.**
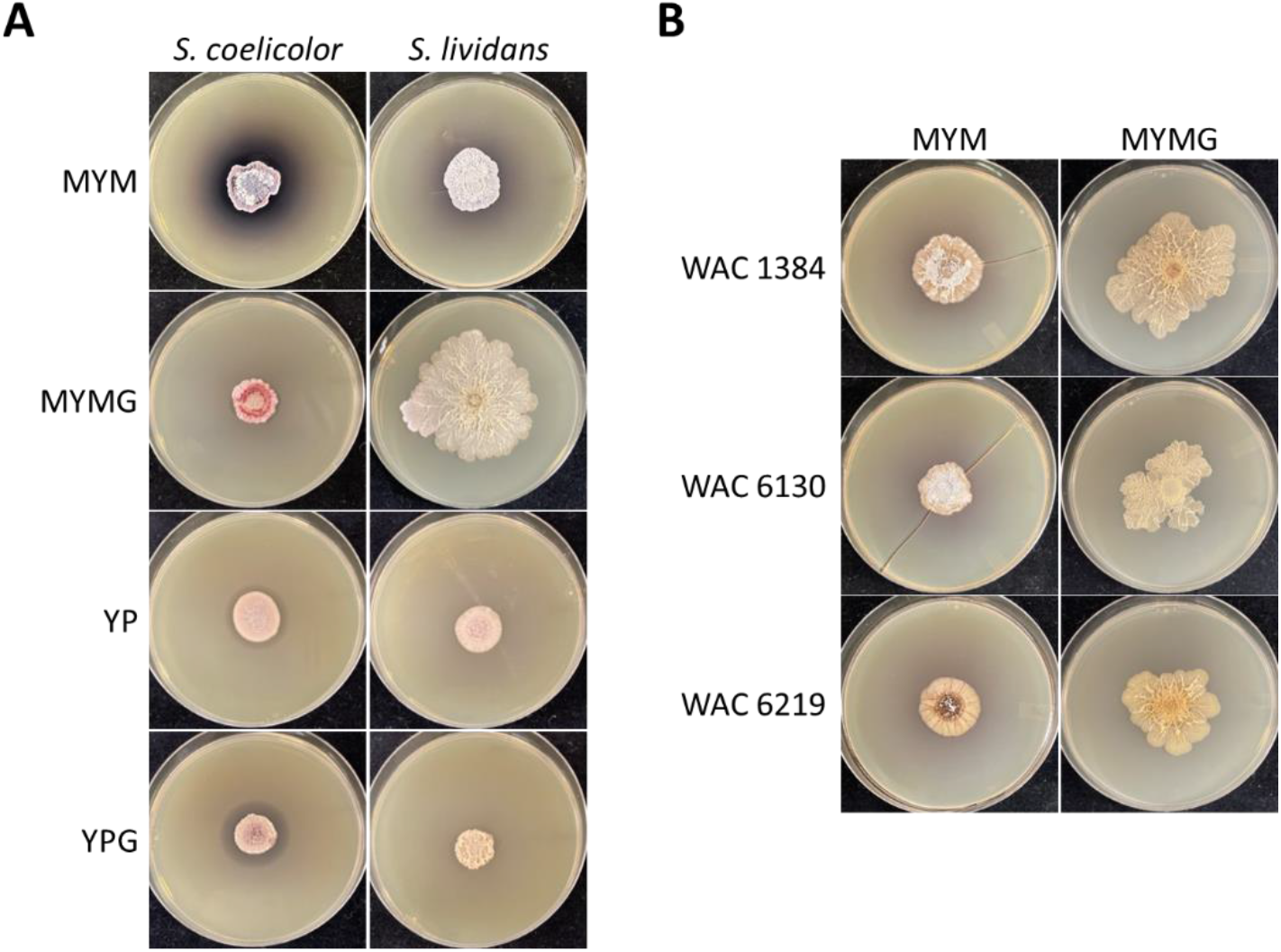
– Diverse *Streptomyces* species display exploratory growth on MYMG. **A)** Growth of the model species *S. coelicolor* and *S. lividans* on MYM, MYMG, YP, and YPG. **B)** Growth of three wild isolates from the Wright Actinomycete Collection on MYM and MYMG. All photos were taken after 14d of growth.

We knew that *sigE* was required for *S. venezuelae* exploration on MYMG. To determine whether the same genetic requirements were conserved in *S. lividans*, we deleted the ortholog of *sigE* (**Supplemental Figure 7**). Consistent with our *S. venezuelae* observations, loss of *sigE* in *S. lividans* resulted in wild type growth on MYM but severely impacted exploration on MYMG.

## DISCUSSION

*Streptomyces* exploratory growth has historically been viewed as proceeding independently of its classical sporulating life cycle (9, 16). Our findings here reveal that exploration can in fact be effectively integrated into the conventional reproductive growth cycle in a nutrient-dependent manner, with either glycerol or complex nitrogen source supplementation promoting both sporulation and exploration.

The discovery that exploratory growth could also proceed without growth medium alkalinization – presumably due to the lack of trimethylamine production – led us to revisit the characteristics that define exploratory growth. When comparing exploration on MYMG to exploration on other media types, three universally shared characteristics were identified. First, exploration involves rapid colony expansion, driven by enhanced growth, coupled with possible surfactant-mediated sliding along the solid growth substrate. Second, exploring colonies adopt a highly wrinkled colony architecture reminiscent of biofilms formed by other bacteria. And finally, exploration requires the function of the sigma factor SigE, which in *Streptomyces* bacteria is responsible for initiating a cell wall stress response (20, 21, 23). There also appear to be common environmental or nutritional factors that impact exploration, including glucose levels (abundant glucose represses exploration), glycerol levels (abundant glycerol enhances exploration), amino acid levels (amino acid abundance influences both sporulation and exploration), and iron levels (multiple iron sequestration strategies are employed by exploring cultures).

Previous work has revealed that trimethylamine production and release by exploring colonies is a powerful mediator of change in the dynamics and composition of the surrounding microbial community (9, 12). In particular, the resulting environmental alkalinization promotes exploratory behaviour in other streptomycetes, and simultaneously inhibits the growth of other microbes by reducing bioavailable iron. The discovery here of an exploration mode that does not involve medium alkalinization suggests there may be situations where such wide-spread environmental modulation is not advantageous. This ‘stealth’ mode of reproductive exploration was not observed under conditions of trimethylamine release, suggesting that these cells may be responsive to different community pressures. We did note, however, that MYMG-grown exploring colonies retained the ability to effectively complete for iron with other microbes on a more local scale through their release of the diffusible foroxymithine siderophore. Foroxymithine can function to reserve iron for its producer strain, and consequently these exploring colonies would be expected to influence iron availability in their immediate vicinity. A future priority will be to probe these different exploration behaviours to better understand their associated fitness costs and benefits.

We had previously assumed that the siderophore requirement of exploring *S. venezuelae* stemmed at least in part from the challenges associated with acquiring iron in an alkaline environment. However, desferrioxamine and foroxymithine siderophores appear to be even more important for *S. venezuelae* growth on MYM and exploration on MYMG (where the pH remains neutral), than during growth under other exploring conditions. An additional surprise was the relative importance of the rare foroxymithine siderophore for the growth of *S. venezuelae,* compared with the near-universal *Streptomyces* siderophore desferrioxamine. These results implied that despite the broad conservation of desferrioxamine biosynthetic potential across the streptomycetes (14, 15), alternative siderophores play key roles in *Streptomyces* growth and development. This is consistent with previous observations in *Streptomyces coelicolor* and *Streptomyces ambofaciens* where it was suggested that either desferrioxamine or the alternative coelichelin siderophore could support colony growth (24). Given the increasing appreciation of the biological and ecological importance of siderophores (25–27), alongside the growing interest in siderophores for their antibiotic potential and their development as antibiotic delivery vehicles (28–32), there is value in better understanding the different properties and relative importance of siderophores in bacteria more generally.

How exploration is genetically regulated remains an important question across all stimulating media conditions. While exploration by developmental mutants on YP and YPG can proceed at varying rates, exploration on MYMG was arrested in Δ*bldD* and Δ*rsiG* strains, likely due to inappropriately high levels of WhiI activity (*whiI* expression is indirectly controlled by both RsiG and BldD (17, 33, 34)). The role of BldD and RsiG as important determinants of MYMG exploration success is particularly interesting as these are two of the three known c-di-GMP-binding protein effectors in *Streptomyces* (the third being the glycogen debranching enzyme GlgX, which impacts energy storage) (17, 18, 35, 36). Nucleotide second messengers like c-di-GMP, have widely been reported to influence population level behaviours like motility and biofilm formation in many bacteria (37–41). Whether nucleotide second messengers contribute to the initiation or continuation of exploratory growth is an area of interest for future studies.

While the requirement of functional BldD and RsiG were unique to MYMG-mediated exploration, the requirement of SigE for successful exploration was consistent across multiple conditions (MYMG, YP and YPG), and multiple *Streptomyces* species. Indeed, the observation of exploratory growth by *S. lividans,* and the conservation of genetic controls across phylogenetically diverse streptomycetes (*S. venezuelae* and *S. lividans*), suggests that exploratory growth is a broadly conserved developmental adaptation in the streptomycetes, and is one that can be triggered by distinct environmental conditions. In the case of *sigE,* its expression is induced in response to cell envelope stress (20–22). These findings suggest that irrespective of the growth condition, entry into exploration may be accompanied by perturbation of the cell wall or membrane. Analogous connections between cell wall integrity and biofilm formation have been reported previously for other microbes, including *Bacillus subtilis* (42), *Staphylococcus aureus* (43), and *Aspergillus fumigatus* (44).

The discovery that *Streptomyces* sporulation and exploration are not mutually exclusive provide interesting bet-hedging opportunities. Exploring colonies move rapidly along solid surfaces and enable colonization of areas distant from their site of inoculation. In contrast, sporulation provides a reservoir of dormant, metabolically inactive cells that can resist many environmental insults. Sporulation by exploring cultures could therefore provide the mobile exploring population with a protected genetic reservoir that would be impervious to nutritional challenges or attack by phage or other microbes, ensuring colony survival as the exploring cells enter new territories. At the same time, the exploring colony offers a novel mechanism of spore dispersal, either directly transferring spores to new environments, or moving them to locations where they may be better dispersed by other factors (e.g. wind, insects).

In other sporulating microbes like *Bacillus subtilis* the decision to sporulate, form a biofilm, or migrate, are not mutually exclusive at a population level, and are increasingly being recognized as outcomes that exist along a regulatory continuum (45, 46), with individual cells or subpopulations having defined – and distinct – roles within the larger population (47). Our results are suggesting that there are shared regulatory elements that contribute to both exploration and sporulation in the streptomycetes, and understanding the interplay between vegetative growth, cell dormancy, biofilm formation, and mobility will be an important goal for the future.

## MATERIALS AND METHODS

### Strains, plasmids, media, and culture conditions

Strains, plasmids, and primers used in this study are listed in Supplemental Tables S3-5, respectively.

*S. venezuelae* NRRL B-65442 was grown in liquid MYM (1% malt extract, 0.4% yeast extract, 0.4% maltose) for overnight cultivation and on solid MYM (2% agar) for spore stock generation. Spore stocks for wild isolates were similarly prepared. Spore stocks for *Streptomyces coelicolor* M145 and *Streptomyces lividans* 1326 were prepared from lawns cultivated on cellophane discs on top of solid MS (2% soya flour, 2% mannitol, 2% agar). When studying growth on solid media, 10 µL of an overnight culture of *S. venezuelae* were spotted onto 40 mL plates containing 2% agar plus additional nutrients [YP (1% yeast extract, 2% peptone), YPG (YP with 2% glycerol), MYM, or MYMG (MYM with 2% glycerol)]. For all other *Streptomyces* species, plates were spotted with 10 µL of a concentrated spore stock. For both solid and liquid media, YP and YPG were prepared using deionized water, while MYM and MYMG were prepared using tap water. Solid MYM or MYMG plates were additionally supplemented with 2% carbon source (e.g., arabinose or galactose) or additional nutrients (sodium nitrate, ammonium chloride [1 µM, 10 µM, 100 µM, or 1 mM]; iron (III) chloride [1 µM, 10 µM, 20 µM, 50 µM, or 100 µM]; alanine, casamino acids, peptone [0.1%, 0.5%, 1%, or 2%]; or 2,2’-dipyridyl [10 µM, 40 µM, 160 µM, or 320 µM]) where appropriate. All *Streptomyces* cultures were grown at 30°C.

For culture-based antibiotic activity assays (detailed below), *Micrococcus luteus* was cultured overnight with shaking at 37 °C in liquid lysogeny broth (LB) (1% tryptone, 0.5% yeast extract, 1% NaCl). Overnight cultures were used to inoculate solid LB medium (1.5% agar; supplemented with 2% glycerol when testing activity of YPG-conditioned medium to eliminate background effects) at a concentration of 1% (vol/vol) and poured as 20 mL plates.

### Construction of *Streptomyces* mutant strains

Gene deletions were generated using ReDirect technology (48). Coding sequences on a cosmid vector carrying large fragments (30 to 40 kb) of genomic DNA from *S. venezuelae* or *S. coelicolor* were replaced by an *oriT*-containing apramycin (Δ*whiH*/Δ*vnz*_*27205*; Δ*whiI*/Δ*vnz*_*28820*; Δ*sigE/*Δ*sli*_3698 in *S. lividans*, using *S. coelicolor* cosmid StE94); or hygromycin (Δ*whiI*/Δ*vnz*_*28820*, or Δ*whiG*/Δ*vnz_26215*) resistance cassette. In creating the *S. venezuelae sigE*/*vnz_15840* deletion strain, the coding sequence and flanking 4-5 kb up– and downstream sequences were PCR amplified and cloned into the pCR2.1-TOPO vector between the HindIII-SpeI and SpeI-XbaI restriction sites (this region lacked an appropriate cosmid for use in creating the gene deletion), before the *sigE* gene was targeted for replacement with the apramycin resistance cassette as described above. The *rsiG* deletion construct was generated through Gibson assembly of four DNA fragments generated by PCR (the vector backbone of pCR2.1-TOPO, a 4.5 kb fragment of *S. venezuelae* genomic DNA upstream of *rsiG*, an *oriT*-containing hygromycin resistance cassette, and a 4.7 kb fragment of *S. venezuelae* genomic DNA downstream of *rsiG*). The mutant cosmids/plasmids were introduced into the non-methylating *Escherichia coli* strain ET12567/pUZ8002, followed by conjugation into *S. venezuelae*. The resulting exconjugants were screened for double-crossover events, and gene deletions were verified by PCR using combinations of primers located upstream, downstream, and internal to the deleted regions (**Supplemental Table 5**).

To generate the *bldM, whiH,* and *whiI* overexpression constructs, a 289-bp fragment containing the *ermE*\* promoter was amplified from pIJ12251 with primers incorporating AvrII and HindIII restriction enzyme recognition sites (**Supplemental Table 5**) and the resulting amplicon was then cloned into pMS82 following digestion with AvrII and HindIII. Coding sequences for *bldM*/*vnz_22005, whiH*/*vnz_27205,* and *whiI/ vnz_28820* were subsequently amplified from *S. venezuelae* genomic DNA, *S. venezuelae* cosmid 4O01, or *S. venezuelae* cosmid Sv-6-D05, respectively with primers that contained 5′ HindIII and KpnI restriction enzyme recognition sites (**Supplemental Table 5**). The resulting amplicons were then cloned into the pMS82-*ermE*\*p construct following digestion with HindIII and KpnI (**Supplemental Table 4**).

To generate the *sigE* and *murA2* overexpression constructs, a 487-bp fragment containing the *ermE*\* promoter was digested out from pIJ12251 using PvuII and EcoRV and subcloned into EcoRV-linearized pMS82. The coding sequence of *sigE* (*vnz_15840*) was amplified from *S. venezuelae* genomic DNA with primers incorporating 5′ NdeI and XhoI restriction enzyme recognition sites and *murA2* (*vnz_28735*) was amplified from *S. venezuelae* genomic DNA with primers incorporating 5′ NdeI and SpeI restriction enzyme recognition sites (**Supplemental Table 5**). The resulting amplicons were then cloned into the pMS82-*ermE*\*p construct following digestion by NdeI and XhoI or NdeI and SpeI (**Supplemental Table 4**).

### Time-lapse videos of exploring colonies

Ten µL of overnight *S. venezuelae* cultures were spotted onto solid medium, and the plates were then placed on an Epson Perfection V800 photo scanner in a 30°C incubator. Images were acquired every hour, and the resulting time course images were compiled in a video format as sequential single frames.

### Growth curves in liquid medium

Wild type *S. venezuelae* grown overnight in 10 mL MYM was used to inoculate 50 mL of fresh MYM or MYMG in baffled flasks to an initial optical density at 600 nm (OD_600_) of 0.025. OD_600_ measurements were taken every hour from 3-10h of growth for one batch of flasks and from 10-20h for a second batch of flasks. Technical triplicates were performed for each growth condition from a single biological replicate.

### Growth curves in solid medium

Photos of growing colonies were taken at set time intervals. Surface area analyses were performed using ImageJ. The scale was established for each image by setting the diameter of the petri dish to 10 cm. The perimeter of the colony was traced, and the area of the captured region was determined. Six replicates were analyzed for each timepoint and condition. For determining changes in colony biomass over time, a single wild type *S. venezuelae* overnight culture (10 mL MYM) was used to inoculate (10 µL) MYM and MYMG plates. Every 2d from 2-12d post-inoculation, whole colonies were scraped from the medium and their wet weight determined using an analytical balance. Three colonies were analyzed for each timepoint and condition.

### RNA isolation, library preparation, and cDNA sequencing

RNA was isolated, as described previously (49), from two independent replicates of *S. venezuelae* grown on solid medium for 2, 4, or 7 days. rRNA was depleted from all samples using a Ribo-Zero rRNA depletion kit. cDNA and Illumina library preparation were achieved using a NEBNext Ultra directional library kit, followed by sequencing using un-paired-end 80-bp reads on the MiSeq Illumina platform. Bioinformatic analyses were conducted using the free open-access platform Galaxy (https://usegalaxy.org). Reads were aligned to the *S. venezuelae* genome using Bowtie2 (50), after which they were sorted, indexed, and converted to BAM format using SAMtools (51). Transcript level normalization and differential transcript level analyses were conducted using DESeq2 (52). DESeq2 transcript normalization and differential gene expression between MYM and MYMG at 4d of growth involved comparing transcripts sequenced from those specific timepoints/conditions as input data; DESeq2 transcript normalization and differential gene expression between MYMG at 2d and 7d of growth in turn involved comparing transcripts sequenced from those specific conditions as well as MYMG at 4d of growth as input data. COG analysis was performed using EggNOG-mapper (53). RNA-seq data has been submitted to the NCBI GEO repository and assigned the accession number GSE240807.

### Culture-based assays for antibiotic activity

LB agar inoculated with our general antibiotic-sensitive indicator organism, *M. luteus*, was prepared as described above. To assay conditioned solid medium, the opening of an inverted sterile 1 mL pipette tip was used to punch an agar plug from the centre of an *S. venezuelae* colony. The agar plug was then transferred to the surface of the indicator agar plate and the resulting plates/plugs incubated overnight at 37 °C. Following incubation, antibiotic activity could be observed as zones of bacterial growth inhibition around the plugs, relative to background growth over the rest of the plate.

### Chrome Azurol S assay for siderophore activity

Samples containing siderophores of interest were assessed using a Chrome Azurol S-based assay as described previously (11). Briefly, sterile solid medium containing the colorimetric dye (2% agar, 100 μM Chrome Azurol S, 200 μM hexadecyltrimethylammonium bromide, 10 μM FeCl_3_, 0.5 M 2-(N-morpholino)ethanesulfonic acid, pH 5.5) was poured as 15 mL plates. As described for the culture-based activity assay, plugs of conditioned media were placed on the surface of the colorimetric indicator plates. Prepared plates were incubated in the dark at room temperature overnight. Following incubation, siderophore activity could be observed as the development of a pink halo around plugs against the blue background of the plate. To measure siderophore activity for conditioned medium *in situ*, 15 mL of CAS agar was prepared and poured as described above into a 10 cm Petri dish. After 7d of growth, biomass was removed from the surface of the conditioned solid medium, followed by overlaying of the conditioned medium with CAS agar. These sandwiched agar plates were then incubated in the dark at room temperature for 4 h, after which the CAS agar layer was removed and photographed.

## Supplemental Video Captions

**Supplemental Video 1:** Wild type *S. venezuelae* grown on MYMG, with images captured every hour for 410 hours.

**Supplemental Video 2:** Wild type *S. lividans* grown on MYM (left) and MYMG (right), with images captured every hour for 410 hours.

**Supplemental Figure 1.**
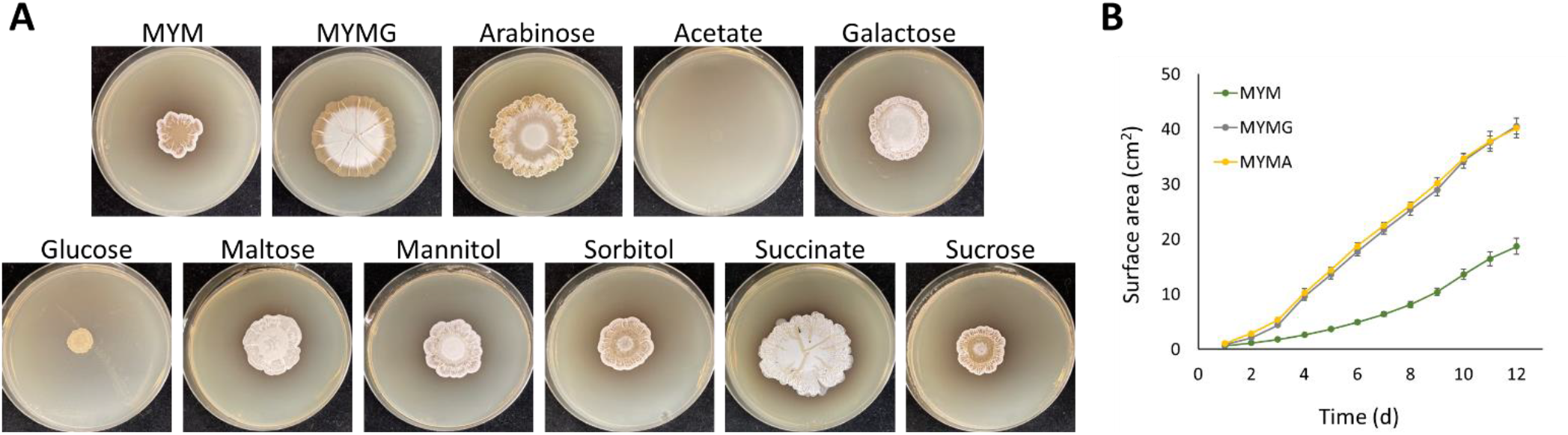
– Effects of other carbon sources on MYM growth. **A** Wild type *S. venezuelae* was spotted to MYM supplemented with different carbon sources at a final concentration of 2% and imaged after 7d of growth. **B)** Solid growth curve measuring surface area expansion over time for wild type *S. venezuelae* spotted to MYM, MYMG, and MYM supplemented with 2% arabinose (MYMA). Error bars represent one standard deviation.

**Supplemental Figure 2.**
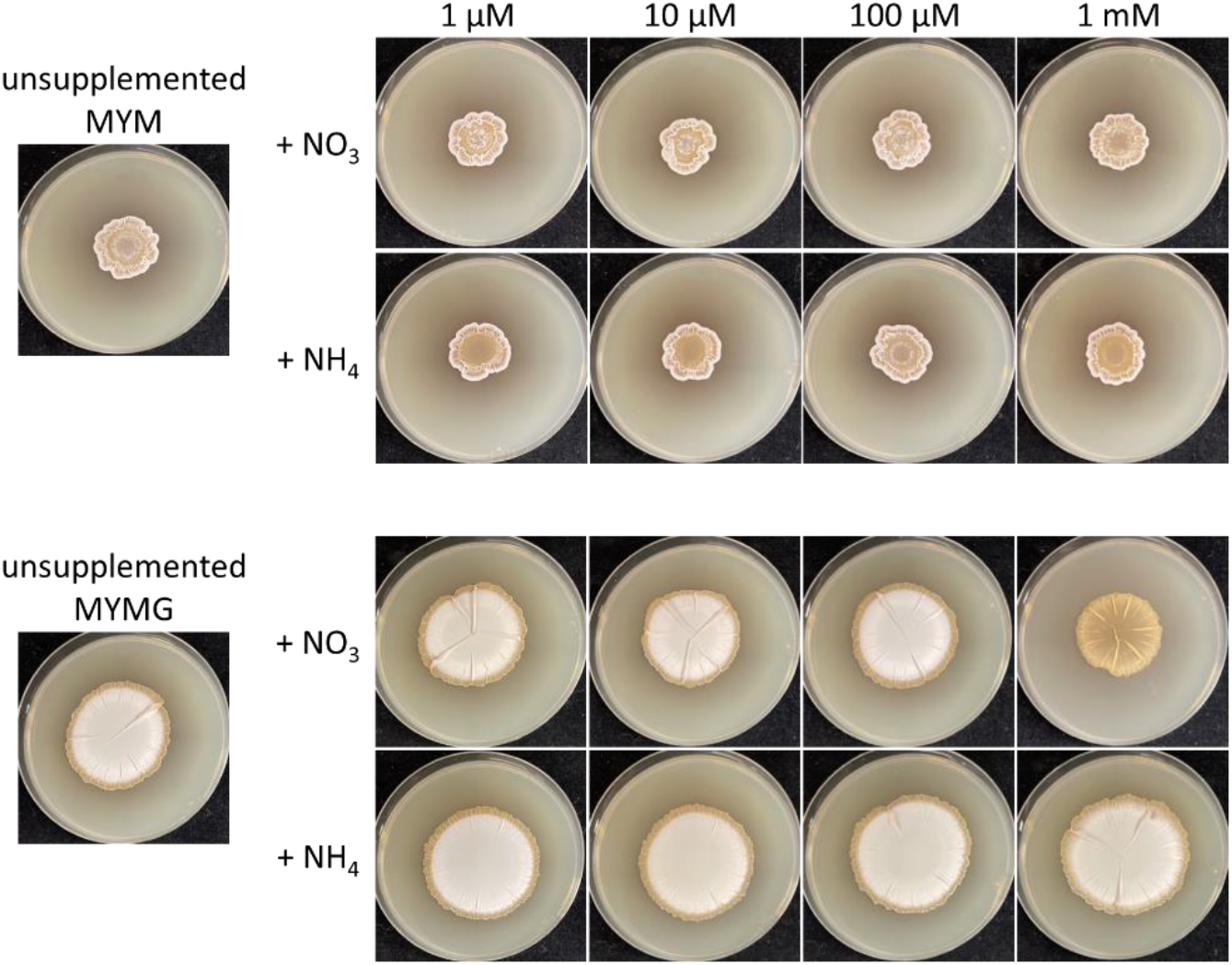
– Effects of inorganic nitrogen supplementation on MYM and MYMG. Wild type *S. venezuelae* was spotted to solid medium supplemented with different concentrations of nitrate or ammonium and photographed after 7d of growth.

**Supplemental Figure 3.**
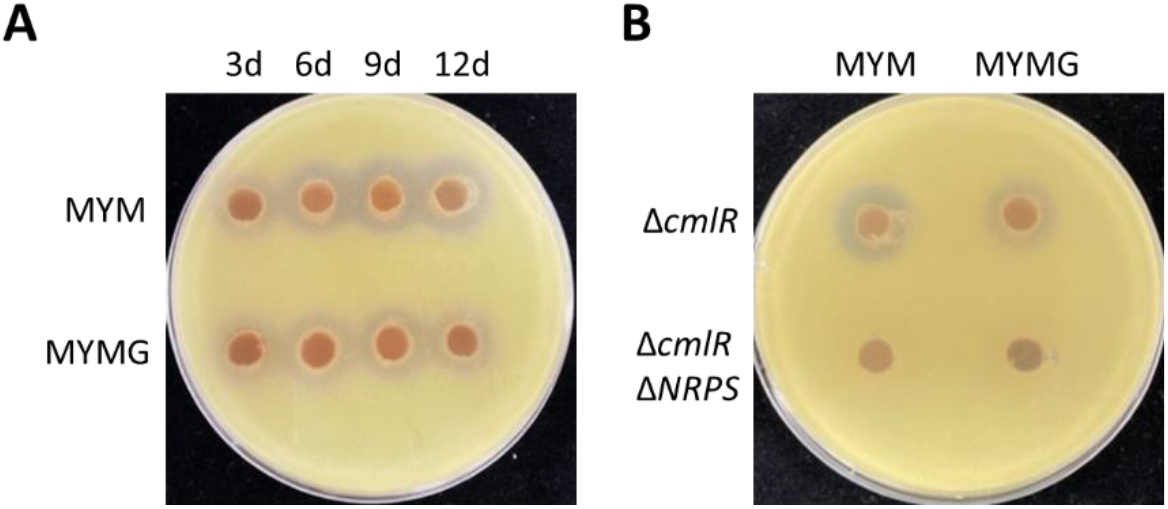
– Antibiotic activity of MYM– and MYMG-grown cultures. **A**) Plugs of conditioned medium from wild type *S. venezuelae* grown for 3-12d on either MYM or MYMG were tested for antibiotic activity against the indicator organism *Micrococcus luteus*. **B)** Antibiotic activity was tested for plugs of medium that had been conditioned for 6d by strains of *S. venezuelae* deficient in chloramphenicol production (Δ*cmlR*) or both chloramphenicol and foroxymithine production (Δ*cmlR* Δ*NRPS*).

**Supplemental Figure 4.**
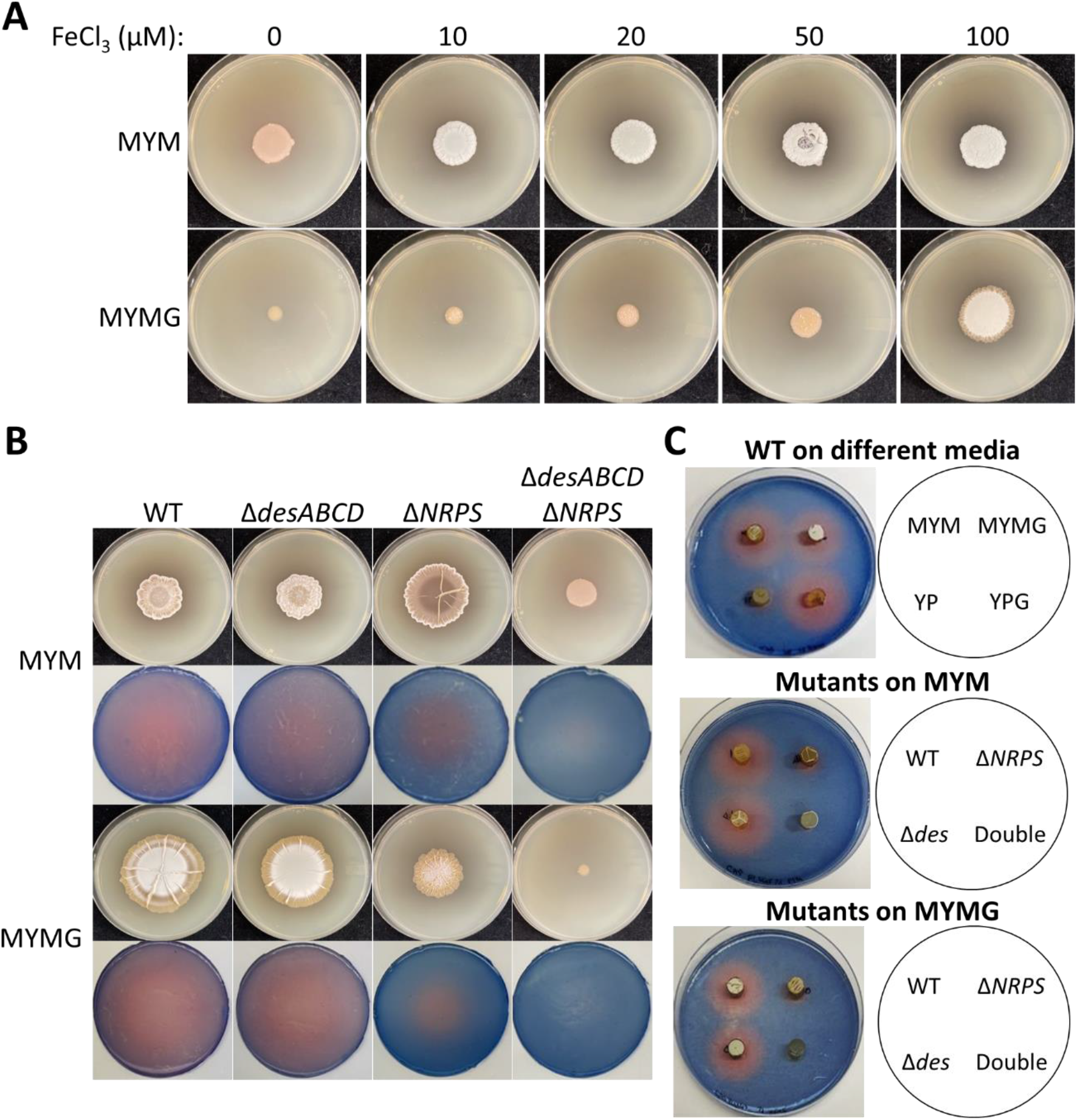
– Effect of iron supplementation on a double siderophore mutant and siderophore diffusibility on MYM and MYMG. **A**) The double siderophore mutant, deficient in both desferrioxamine and foroxymithine production (Δ*desABCD* Δ*NRPS*), was spotted to MYM and MYMG supplemented with a broader concentration range of FeCl_3_ than tested in **Figure 4A**. Images were taken after 7d of growth. **B)** A panel of siderophore mutant strains were spotted to MYM and MYMG and imaged after 7d of growth (first and third rows). CAS agar applied to the resulting conditioned medium was developed for 4h before being removed and imaged (second and fourth rows) resulting in zones of colour change local to where siderophores were active in the conditioned medium. **C)** Plugs of conditioned media from 7d colonies were transferred onto CAS agar. Following overnight incubation, pink zones were observed indicating diffusible siderophore activity originating from the conditioned medium.

**Supplemental Figure 5.**
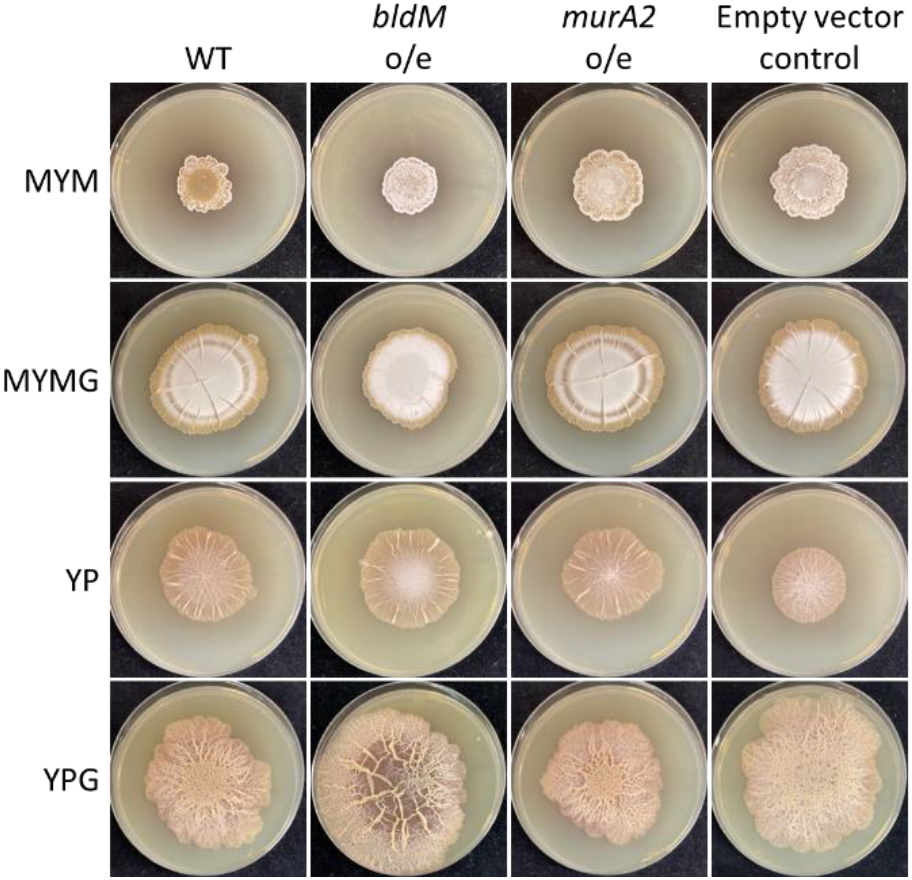
– Overexpressing *bldM* or *murA2* fails to recapitulate a *whiI* overexpression phenotype. Coding sequences for *bldM* and *murA2* were cloned under the control of the strong constitutive *ermE*\* promoter, and the resulting constructs were introduced into the wild type and then spotted to different media. A wild type control strain carrying the integrating plasmid with *ermE*\* alone was included for comparison (right column). Images were taken after 7d of growth.

**Supplemental Figure 6.**
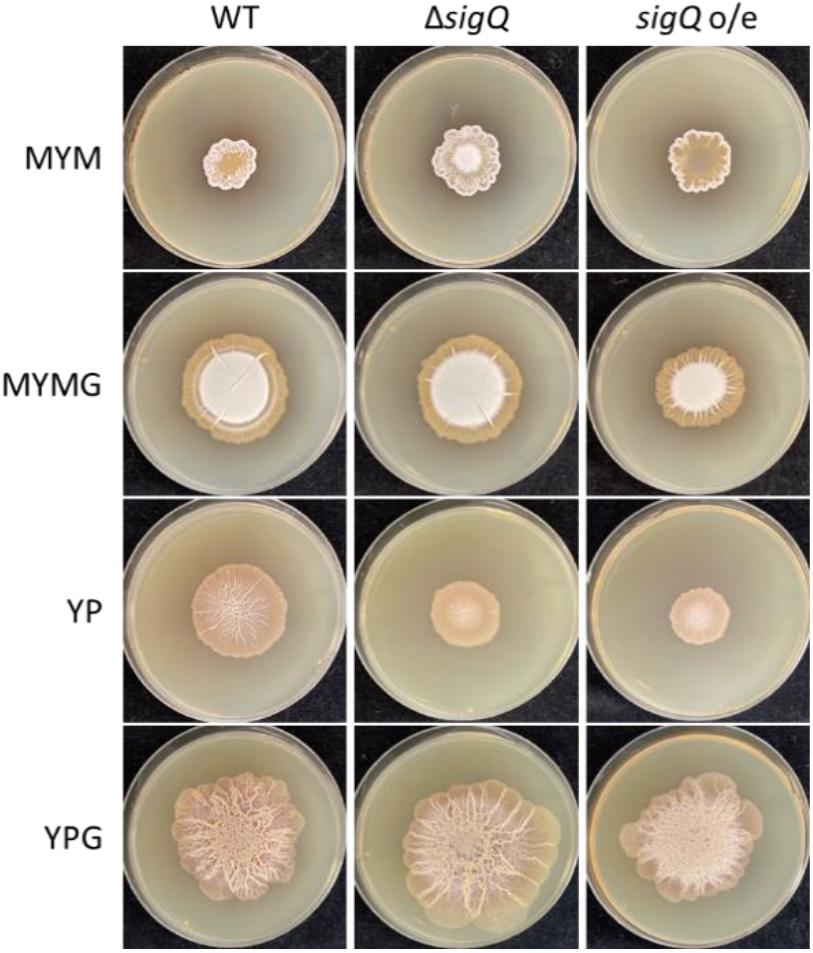
– Contribution of *sigQ* to *S. venezuelae* development. *S. venezuelae* strains carrying a deletion in the *sigQ* or a plasmid with *sigQ* under the strong constitutive promoter *ermE*\* were spotted to different media and photographed after 7d of growth.

**Supplemental Figure 7.**
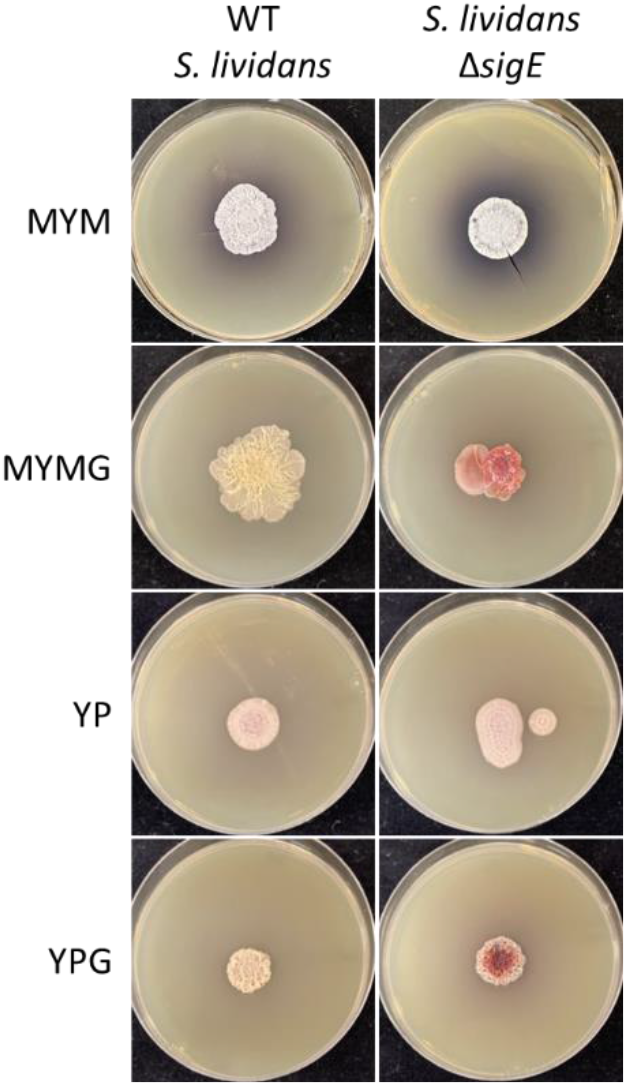
– SigE contributes to MYMG exploration in *S. lividans*. The homolog of *sigE* was deleted in *S. lividans* and spotted alongside the wild type strain on different media. Photos were taken at 14d of growth.

## REFERENCES

1. J. Bérdy, Bioactive microbial metabolites. J. Antibiot. 58, 1–26 (2005).

2. N. Tsuji, M. Kobayashi, K. Nagashima, Y. Wakisaka, K. Koizumi, A new antifungal antibiotic, trichostatin. J. Antibiot. 29, 1–6 (1976).

3. T. Kaur, A. Vasudev, S. K. Sohal, R. K. Manhas, Insecticidal and growth inhibitory potential of *Streptomyces hydrogenans* DH16 on major pest of India, Spodoptera litura (Fab.) (Lepidoptera: Noctuidae). BMC Microbiol. 14 (2014).

4. G. Hoerlein, “Glufosinate (phosphinothricin), a natural amino acid with unexpected herbicidal properties.” in Reviews of Environmental Contamination and Toxicology, G. W. Ware, Ed. (Springer, 1994), pp. 73–145.

5. A. Bolourian, Z. Mojtahedi, Immunosuppressants produced by *Streptomyces*: evolution, hygiene hypothesis, tumour rapalog resistance and probiotics. Environ. Microbiol. Rep. 10, 123–126 (2018).

6. C. Reading, M. Cole, Clavulanic acid: a beta-lactamase-inhibiting beta-lactam from *Streptomyces clavuligerus*. Antimicrob. Agents Chemother. 11, 852–7 (1977).

7. M. J. Buttner, K. Flärdh, M. Howard, A. M. Hempel, D. M. Richards, Regulation of apical growth and hyphal branching in *Streptomyces*. Curr. Opin. Microbiol. 15, 737–743 (2012).

8. M. P. Zambri, M. A. Williams, M. A. Elliot, “How Streptomyces thrive: Advancing our understanding of classical development and uncovering new behaviors” in Advances in Microbial Physiology, Advances in Microbial Physiology., R. K. Poole, D. J. Kelly, Eds. (Academic Press, 2022), pp. 203–236. (2022)

9. S. E. Jones, et al., *Streptomyces* exploration is triggered by fungal interactions and volatile signals. eLife 6, e21738 (2017).

10. E. M. F. Shepherdson, T. Netzker, Y. Stoyanov, M. A. Elliot, Exploratory growth in *Streptomyces venezuelae* involves a unique transcriptional program, enhanced oxidative stress response, and profound acceleration in response to glycerol. J. Bacteriol. 204, e00623–21 (2022).

11. E. M. F. Shepherdson, M. A. Elliot, Cryptic specialized metabolites drive Streptomyces exploration and provide a competitive advantage during growth with other microbes. Proc. Natl. Acad. Sci. 119, e2211052119 (2022).

12. S. E. Jones, et al., *Streptomyces* volatile compounds influence exploration and microbial community dynamics by altering iron availability. mBio 10, e00171–19 (2019).

13. X. Zhang, S. N. Andres, M. A. Elliot, Interplay between nucleoid-associated proteins and transcription factors in controlling specialized metabolism in Streptomyces. mBio 12, e01077–21 (2021).

14. M. Imbert, M. Béchet, R. Blondeau, Comparison of the main siderophores produced by some species of *Streptomyces*. Curr. Microbiol. 31, 129–133 (1995).

15. F. Barona-Gómez, U. Wong, A. E. Giannakopulos, P. J. Derrick, G. L. Challis, Identification of a cluster of genes that directs desferrioxamine biosynthesis in *Streptomyces coelicolor* M145. J. Am. Chem. Soc. 31, 129–133 (2004).

16. S. E. Jones, M. A. Elliot, ‘Exploring’ the regulation of *Streptomyces* growth and development. Curr. Opin. Microbiol. 42, 25–30 (2018).

17. K. A. Gallagher, et al., c-di-GMP arms an anti-σ to control progression of multicellular differentiation in *Streptomyces*. Mol. Cell 77, 586–599 (2020).

18. M. A. Schumacher, et al., Evolution of a σ–(c-di-GMP)–anti-σ switch. Proc. Natl. Acad. Sci. U. S. A. 118, e2105447118 (2021).

19. M. M. Al-Bassam, M. J. Bibb, M. J. Bush, G. Chandra, M. J. Buttner, Response regulator heterodimer formation controls a key stage in *Streptomyces* development. PLoS Genet. 10, e1004554 (2014).

20. M. S. B. Paget, L. Chamberlin, A. Atrih, S. J. Foster, M. J. Buttner, Evidence that the extracytoplasmic function sigma factor σ(E) is required for normal cell wall structure in Streptomyces coelicolor A3(2). J. Bacteriol. 181, 204–211 (1999).

21. M. S. B. Paget, E. Leibovitz, M. J. Buttner, A putative two-component signal transduction system regulates σ(E), a sigma factor required for normal cell wall integrity in *Streptomyces coelicolor* A3(2). Mol. Microbiol. 33, 97–107 (1999).

22. H. J. Hong, M. S. B. Paget, M. J. Buttner, A signal transduction system in *Streptomyces coelicolor* that activates the expression of a putative cell wall glycan operon in response to vancomycin and other cell wall-specific antibiotics. Mol. Microbiol. 44, 1199–1211 (2002).

23. N. T. Tran, et al., Defining the regulon of genes controlled by σE, a key regulator of the cell envelope stress response in *Streptomyces coelicolor*. Mol. Microbiol. 112, 461–481 (2019).

24. F. Barona-Gómez, et al., Multiple biosynthetic and uptake systems mediate siderophore–dependent iron acquisition in *Streptomyces coelicolor* A3(2) and *Streptomyces ambofaciens* ATCC 23877. Microbiology 152, 3355–3366 (2006).

25. J. Kramer, Ö. Özkaya, R. Kümmerli, Bacterial siderophores in community and host interactions. Nat. Rev. Microbiol. 18, 152–163 (2020).

26. G. L. Challis, D. A. Hopwood, Synergy and contingency as driving forces for the evolution of multiple secondary metabolite production by *Streptomyces* species. Proc. Natl. Acad. Sci. U. S. A. 100, 14555–14561 (2003).

27. M. F. Traxler, M. R. Seyedsayamdost, J. Clardy, R. Kolter, Interspecies modulation of bacterial development through iron competition and siderophore piracy. Mol. Microbiol. 86, 628–644 (2012).

28. G. Swayambhu, M. Bruno, A. M. Gulick, B. A. Pfeifer, Siderophore natural products as pharmaceutical agents. Curr. Opin. Biotechnol. 69, 242–251 (2021).

29. F. Zhang, et al., Thalassosamide, a siderophore discovered from the marine-derived bacterium Thalassospira profundimaris. J. Nat. Prod. 80, 2551–2555 (2017).

30. Y. Takehana, et al., Fradiamine A, a new siderophore from the deep-sea actinomycete *Streptomyces fradiae* MM456M-mF7. J. Antibiot. 70, 611–615 (2017).

31. M. Ghosh, et al., Targeted antibiotic delivery: selective siderophore conjugation with daptomycin confers potent activity against multidrug resistant *Acinetobacter baumannii* both *in vitro* and *in vivo*. J. Med. Chem. 60, 4577–4583 (2017).

32. M. Sassone-Corsi, et al., Siderophore-based immunization strategy to inhibit growth of enteric pathogens. Proc. Natl. Acad. Sci. U. S. A. 113, 13462–13467 (2016).

33. M. A. Elliot, M. J. Bibb, M. J. Buttner, B. K. Leskiw, BldD is a direct regulator of key developmental genes in *Streptomyces coelicolor* A3(2). Mol. Microbiol. 40, 257–269 (2001).

34. C. D. de. Hengst, et al., Genes essential for morphological development and antibiotic production in *Streptomyces coelicolor* are targets of BldD during vegetative growth. Mol. Microbiol. 78, 361–379 (2010).

35. N. Tschowri, et al., Tetrameric c-di-GMP mediates effective transcription factor dimerization to control *Streptomyces* development. Cell 158, 1136–1147 (2014).

36. M. A. Schumacher, et al., Allosteric regulation of glycogen breakdown by the second messenger cyclic di-GMP. Nat. Commun. 13, 1–14 (2022).

37. D. Kalia, et al., Nucleotide, c-di-GMP, c-di-AMP, cGMP, cAMP, (p)ppGpp signaling in bacteria and implications in pathogenesis. Chem. Soc. Rev. 42, 305–341 (2013).

38. M. Merighi, V. T. Lee, M. Hyodo, Y. Hayakawa, S. Lory, The second messenger bis-(3′-5′)– cyclic-GMP and its PilZ domain-containing receptor Alg44 are required for alginate biosynthesis in *Pseudomonas aeruginosa*. Mol. Microbiol. 65, 876–895 (2007).

39. C. Baraquet, K. Murakami, M. R. Parsek, C. S. Harwood, The FleQ protein from *Pseudomonas aeruginosa* functions as both a repressor and an activator to control gene expression from the *pel* operon promoter in response to c-di-GMP. Nucleic Acids Res. 40, 7207–7218 (2012).

40. P. V. Krasteva, et al., *Vibrio cholerae* Vpst regulates matrix production and motility by directly sensing cyclic di-GMP. Science. 327, 866–868 (2010).

41. D. Srivastava, M. L. Hsieh, A. Khataokar, M. B. Neiditch, C. M. Waters, Cyclic di-GMP inhibits *Vibrio cholerae* motility by repressing induction of transcription and inducing extracellular polysaccharide production. Mol. Microbiol. 90, 1262–1276 (2013).

42. T. Bucher, Y. Oppenheimer-Shaanan, A. Savidor, Z. Bloom-Ackermann, I. Kolodkin-Gal, Disturbance of the bacterial cell wall specifically interferes with biofilm formation. Environ. Microbiol. Rep. 7, 990–1004 (2015).

43. S. Dubrac, I. G. Boneca, O. Poupel, T. Msadek, New insights into the WalK/WalR (YycG/YycF) essential signal transduction pathway reveal a major role in controlling cell wall metabolism and biofilm formation in *Staphylococcus aureus*. J. Bacteriol. 189, 8257– 8269 (2007).

44. Y. Chen, et al., The transcription factor SomA synchronously regulates biofilm formation and cell wall homeostasis in *Aspergillus fumigatus*. mBio 11, e02329–20 (2020).

45. R. R. Grau, et al., A duo of potassium-responsive histidine kinases govern the multicellular destiny of *Bacillus subtilis*. mBio 6, e00581–15 (2015).

46. Z. Chen, B. Zarazúa-Osorio, P. Srivastava, M. Fujita, O. A. Igoshin, The slowdown of growth rate controls the single-cell distribution of biofilm matrix production via an SinI-SinR-SlrR network. mSystems 8, 1–19 (2023).

47. S. M. Yannarell, et al., Extensive cellular multi-tasking within *Bacillus subtilis* biofilms. mSystems, e00891–22 (2023).

48. B. Gust, G. L. Challis, K. Fowler, T. Kieser, K. F. Chater, PCR-targeted *Streptomyces* gene replacement identifies a protein domain needed for biosynthesis of the sesquiterpene soil odor geosmin. Proc. Natl. Acad. Sci. U. S. A. 100, 1541–1546 (2003).

49. M. J. Moody, R. A. Young, S. E. Jones, M. A. Elliot, Comparative analysis of non-coding RNAs in the antibiotic-producing *Streptomyces* bacteria. BMC Genomics 14, 1–17 (2013).

50. B. Langmead, S. L. Salzberg, Fast gapped-read alignment with Bowtie 2. Nat. Methods 9, 357–359 (2012).

51. H. Li, et al., The Sequence Alignment/Map format and SAMtools. Bioinformatics 25, 2078–9 (2009).

52. M. I. Love, W. Huber, S. Anders, Moderated estimation of fold change and dispersion for RNA-seq data with DESeq2. Genome Biol. 15, 1–21 (2014).

53. C. P. Cantalapiedra, A. Hernández-Plaza, I. Letunic, P. Bork, J. Huerta-Cepas, eggNOG-mapper v2: Functional annotation, orthology assignments, and domain prediction at the metagenomic scale. Mol. Biol. Evol. (2021) 10.1093/molbev/msab293.

