## Supplementary Tables 3-5 for "Redefining development in *Streptomyces* bacteria: integrating exploration into the classical sporulating life cycle"

**Supplemental Table 3 – Strains used in this study**

| **Strains** | **Genotype/characteristics/use** | **Reference** |
|---|---|---|
| ***Streptomyces*** | | |
| *S. venezuelae* NRRL B-65442 | Wild type | (1) |
| *S. coelicolor* A3(2) M145 | Wild type | (2) |
| *S. lividans* 1326 | Wild type | Gift from the JIC |
| WAC 1384 | Wild *Streptomyces* isolate | Gift from G. Wright |
| WAC 6130 | Wild *Streptomyces* isolate | Gift from G. Wright |
| WAC 6219 | Wild *Streptomyces* isolate | Gift from G. Wright |
| SV13 | *S. venezuelae bldM::aac(3)IV* (*vnz_22005*) | (3) |
| SV77 | *S. venezuelae bldD::aac(3)IV* (*vnz_05285*) | (4) |
| E331 | *S. venezuelae desA-D::aac(3)IV* | (5) |
| E351 | *S. venezuelae gylR-D::aac(3)IV* (glycerol catabolism operon; *vnz_06115-30*) | (6) |
| E352 | *S. venezuelae sigQ::aac(3)IV* (*vnz_22610*) | (6) and this work |
| E332 | *S. venezuelae cmlR::aac(3)IV* (*vnz_04400*) | (7) |
| E333 | *S. venezuelae vnz_34785::hyg* (foroxymithine non-ribosomal peptide synthetase; *fxmE*) | (8) |
| E334 | *S. venezuelae cmlR::aac(3)IV vnz_34785::hyg* | (8) |
| E335 | *S. venezuelae desA-D::aac(3)IV vnz_34785::hyg* | (8) |
| E353 | *S. venezuelae whiG::hyg* (*vnz_26215*) | This work |
| E354 | *S. venezuelae whiH::aac(3)IV* (*vnz_27205*) | This work |
| E355 | *S. venezuelae whiI::aac(3)IV* (*vnz_28820*) | This work |
| E356 | *S. venezuelae whiI::hyg* | This work |
| E357 | *S. venezuelae rsiG::hyg* (*vnz_19430*) | This work |

**Formatted:** Spanish (Spain)

**Formatted:** Spanish (Spain)

| | | |
|---|---|---|
| E358 | *S. venezuelae rsiG::hyg whiH::aac(3)IV* | This work |
| E359 | *S. venezuelae rsiG::hyg whiI::aac(3)IV* | This work |
| E360 | *S. venezuelae sigE::aac(3)IV* (*vnz_15840*) | This work |
| E361 | *S. venezuelae sigE::aac(3)IV whiI::hyg* | This work |
| E362 | *S. lividans sigE::aac(3)IV* (*sli_3698*) | This work |
| ***Escherichia coli*** | | |
| DH5α | Routine cloning | Invitrogen |
| BW25113/pIJ790 | Introducing mutations in cosmid DNA | (9) |
| ET12567/pUZ8002 | Generation of methylation-free plasmid DNA and conjugation into *Streptomyces* | (9) |
| **Other organisms** | | |
| *Micrococcus luteus* | Antibiotic-sensitive indicator bacterium | Gift from J. Nodwell |

**Formatted:** Spanish (Spain)

**Formatted:** Spanish (Spain)

**Supplementary Table 4 – Plasmids and cosmids used in this study**

| Cosmid/plasmid | Description | Reference |
|---|---|---|
| 4O01 | *S. venezuelae* cosmid carrying *whiH* | Gift from M. Buttner |
| StE94 | *S. coelicolor* cosmid carrying *sigE* | Gift from M. Buttner |
| Sv-2_B03 | *S. venezuelae* cosmid carrying *whiG* | Gift from M. Buttner |
| Sv-6-D05 | *S. venezuelae* cosmid carrying *whiI* | Gift from M. Buttner |
| pIJ773 | Plasmid carrying the *aac(3)IV-oriT* cassette | (9) |
| pIJ10700 | Plasmid carrying the *hyg-oriT* cassette | (10) |
| pIJ12551 | Plasmid carrying the strong *Streptomyces* promoter *ermE**p | (11) |
| pMS82 | Integrative cloning vector: *hyg*, *oriT*, *int* ΦBT1, *attP* ΦBT1 | (12) |
| pMC302 | *ermE**p-*vnz_22610* (*sigQ*) cloned into pMS82 | (6) |
| pMC303 | *ermE**p-*vnz_22005* (*bldM*) cloned into pMS82 | This work |
| pMC304 | *ermE**p-*vnz_28735* (*murA2*) cloned into pMS82 | This work |
| pMC305 | *ermE**p-*vnz_15840* (*sigE*) cloned into pMS82 | This work |
| pMC306 | *ermE**p-*vnz_27205* (*whiH*) cloned into pMS82 | This work |
| pMC307 | *ermE**p-*vnz_28820* (*whiI*) cloned into pMS82 | This work |
| pCR2.1-TOPO | Cloning vector used for creating *sigE* and *rsiG* deletion strains | Invitrogen |
| pMC308 | *sigE,* together with 4-5 kb upstream and downstream flanking sequences cloned into the pCR2.1-TOPO vector | This work |
| pMC309 | 4-5 kb sequences upstream and downstream of the *rsiG* coding sequence cloned into the pCR2.1-TOPO vector separated by a *hyg-oriT* cassette | This work |

**Supplementary Table 5 – Oligonucleotides used in this study**

| **Name** | **Sequence (5' to 3')** | **Use** |
|---|---|---|
| *whiG* ReD FWD | CGACACACCCGACAGCAGAACGGCTCAAGGCAA CGCATG***ATTCCGGGGATCCGTCGACC*** | Creation and confirmation of the Δ*whiG* mutation |
| *whiG* ReD REV | TGGGCACGGCTCCACTGTACGACCGGCTCGCCC CGGTCA***TGTAGGCTGGAGCTGCTTC*** | Creation and confirmation of the Δ*whiG* mutation |
| *whiG* Up FWD | CCCGGGTTCGAAGATGTGGC | Confirmation of the Δ*whiG* mutation |
| *whiG* In REV | GGGTGATGGCGTACGTCTCG | Confirmation of the Δ*whiG* mutation |
| *whiH* ReD FWD | GCCGACAAAGGATGCGTGAGTACCCTTGCGCAC ACCATG***ATTCCGGGGATCCGTCGACC*** | Creation and confirmation of the Δ*whiH* mutation |
| *whiH* ReD REV | GCCCCCGACTCCGCACTCCCGGCCACGGCCCGC GGATCA***TGTAGGCTGGAGCTGCTTC*** | Creation and confirmation of the Δ*whiH* mutation |
| *whiH* In FWD | GGATGCTGGAGCACCTCTCCG | Confirmation of the Δ*whiH* mutation |
| *whiH* Down REV | AGGGTACATTCACGCCGCCC | Confirmation of the Δ*whiH* mutation |
| *whiI* ReD FWD | CGGCTCCGTCCCGCACCCTTCCCCCCAGGAGGCC TGGTG***ATTCCGGGGATCCGTCGACC*** | Creation and confirmation of the Δ*whiI* mutation |
| *whiI* ReD REV | CCGTCGACAGGCGCCGGCTTTCCGTCGGCCGGT CCGTCA***TGTAGGCTGGAGCTGCTTC*** | Creation and confirmation of the Δ*whiI* mutation |
| *whiI* Up FWD | CCTGGTGCGATCTGCTCTGTC | Confirmation of the Δ*whiI* mutation |
| *whiI* In REV | CGATCACGTCGCGTACTCCG | Confirmation of the Δ*whiI* mutation |
| livid *sigE* ReD FWD | CCACCGTCGGAGTACGGGGATCGGAAGGCGGT TGACATG***ATTCCGGGGATCCGTCGACC*** | Creation and confirmation of the *S. lividans* Δ*sigE* mutation |
| livid *sigE* ReD REV | TGGTTCTCCGGGTCCCCCTGGGTGTCCGACCGTC GGTCA***TGTAGGCTGGAGCTGCTTC*** | Creation and confirmation of the *S. lividans* Δ*sigE* mutation |
| livid *sigE* Up FWD | CCGTGACGGACAAATCGCCTC | Confirmation of the *S. lividans* Δ*sigE* mutation |
| livid *sigE* In REV | GCTCGGTCGGCACCTCCTC | Confirmation of the *S. lividans* Δ*sigE* mutation |
| livid *sigE* Down REV | CTGCGTGGTTCTCCGGGTC | Confirmation of the *S. lividans* Δ*sigE* mutation |
| *sigE* TOPO Left arm FWD HindIII | CATCATAAGCTTGCTTCGTACGCTTCGCGGAG | Creation of the *S. venezuelae sigE* cosmid equivalent |
| *sigE* TOPO Left arm REV SpeI | TACTACACTAGTCATGGGCATTCGGACCGTCC | Creation of the *S. venezuelae sigE* cosmid equivalent |
| *sigE* TOPO Right arm FWD SpeI | CATCATACTAGTCGCCCCGCTGTTACACAACC | Creation of the *S. venezuelae sigE* cosmid equivalent |
| *sigE* TOPO Right arm REV XbaI | CATCATTCTAGAGAGGGCCGCTTTGGTGACG | Creation of the *S. venezuelae sigE* cosmid equivalent |
| *sigE* ReD FWD | GAACGAAGCAGGTCACGGGGTTCGGAGGCGGT TCGGATG***ATTCCGGGGATCCGTCGACC*** | Creation and confirmation of the *S. venezuelae* Δ*sigE* mutation |

| | | |
|---|---|---|
| *sigE* ReD REV | TGCTGCCGTCGGCTTCGCCGTCTAGGCCGCGCACCGCTC***TGTAGGCTGGAGCTGCTTC*** | Creation and confirmation of the *S. venezuelae* Δ*sigE* mutation |
| *sigE* Up FWD | GTGCGTCCACCGACGGCTG | Confirmation of the *S. venezuelae* Δ*sigE* mutation |
| *sigE* In FWD | GACGCCTCCGTCGACGACC | Confirmation of the *S. venezuelae* Δ*sigE* mutation |
| *sigE* Down REV | CGGCCATGGACAGCCCGAC | Confirmation of the *S. venezuelae* Δ*sigE* mutation |
| *rsiG* Gibson Up FWD | CTATGGAAAAACGCCAGCAACGCGGCCTTTTTACGGTTCCCGCTGAGCAAGCTCCTGCTG | Creation and confirmation of the Δ*rsiG* mutation |
| *rsiG* Gibson Up REV | AATAGGAACTTCGAACTGCAGGTCGACGGATCCCCGGAATCAGATTCGTCCCCTCGACCG | Creation and confirmation of the Δ*rsiG* mutation |
| *rsiG* Gibson HygOriT FWD | CGCCTCCGCACCGGTCGAGGGGACGAATCTGATTCCGGGGATCCGTCGACC | Creation and confirmation of the Δ*rsiG* mutation |
| *rsiG* Gibson HygOriT REV | CGTCAGGCGAGCAGGTCGTCGACCTGGGTGTAGGCTGGAGCTGCTTCG | Creation and confirmation of the Δ*rsiG* mutation |
| *rsiG* Gibson Down FWD | TCTAGAAAGTATAGGAACTTCGAAGCAGCTCCAGCCTACACCCAGGTCGACGACCTGCTC | Creation and confirmation of the Δ*rsiG* mutation |
| *rsiG* Gibson Down REV | ATTAAGTTGGGTAACGCCAGGGTTTTCCCAGTCACGACGTGAGCACGGCCACGATCTGC | Creation and confirmation of the Δ*rsiG* mutation |
| *rsiG* Gibson TOPO FWD | GTGGCGCGGCTCGGCGCCGGGCAGATCGTGGCCGTGCTCACGTCGTGACTGGGAAAACCC | Creation and confirmation of the Δ*rsiG* mutation |
| *rsiG* Gibson TOPO REV | ACGTAACCCTCCGGTACGGGCAGCAGGAGCTTGCTCAGCGGGAACCGTAAAAAGGCCGCG | Creation and confirmation of the Δ*rsiG* mutation |
| *rsiG* Up FWD | GTGCAGGAGTGGCACCATGC | Confirmation of the Δ*rsiG* mutation |
| *rsiG* In REV | AGGTCCGACAGCTCCACCTC | Confirmation of the Δ*rsiG* mutation |
| *ermE* FWD AvrII | ATATCCTAGGAGCCCGACCCGAGCACGC | Cloning *ermE**p into pMS82 |
| *ermE* REV HindIII | ATATAAGCTTGATCCTACCAACCGGCACGA | Cloning *ermE**p into pMS82 |
| *whiH* OE FWD HindIII | CATCATAAGCTTCAAGGGTCACGGAAGCAACGC | Creation of *whiH* overexpression construct |
| *whiH* OE REV KpnI | CATCATGGTACCCGCCCAATCTTCGTGGCCC | Creation of *whiH* overexpression construct |
| *whiI* OE FWD HindIII | CATCATAAGCTTTCACAGCACGGGTTCGCC | Creation of *whiI* overexpression construct |
| *whiI* OE REV KpnI | CATCATGGTACCCCAGCAACGGGATCGGAAGC | Creation of *whiI* overexpression construct |
| *bldM* OE FWD HindIII | CATCATAAGCTTGCTACAGCTGCGAGAGCCGCG | Creation of *bldM* overexpression construct |
| *bldM* OE REV KpnI | CATCTTGGTACCCTCCATGGTGCCCCGACGCC | Creation of *bldM* overexpression construct |
| *sigE* OE FWD NdeI | CGAGTCCATATGAGGAACGAAGCAGGTCACGGG | Creation of *sigE* overexpression construct |
| *sigE* OE REV XhoI | CATCATCTCGAGCGTCGGCTTCGCCGTCTAG | Creation of *sigE* overexpression construct |

| | | |
|---|---|---|
| *murA2* OE FWD NdeI | CATCATCATATGCCTACCCCAACTGCGAGTG | Creation of *murA2* overexpression construct |
| *murA2* OE REV SpeI | CATCATACTAGTGGAGGCCTGGGACTCGATGAC | Creation of *murA2* overexpression construct |

Cassette-specific sequence has been bolded and italicized. Restriction enzyme recognition sequences are underlined.


1. Gomez-Escribano JP, Holmes NA, Schlimpert S, Bibb MJ, Chandra G, Wilkinson B, Buttner MJ, Bibb MJ. 2021. *Streptomyces venezuelae* NRRL B-65442: genome sequence of a model strain used to study morphological differentiation in filamentous actinobacteria . J Ind Microbiol Biotechnol.

2. Kieser T, Bibb MJ, Buttner MJ, Chater KF. 2000. Practical *Streptomyces* Genetics. John Innes Foundation.

3. Al-Bassam MM, Bibb MJ, Bush MJ, Chandra G, Buttner MJ. 2014. Response regulator heterodimer formation controls a key stage in *Streptomyces* development. PLoS Genet 10:e1004554.

4. Tschowri N, Schumacher MA, Schlimpert S, Chinnam NB, Findlay KC, Brennan RG, Buttner MJ. 2014. Tetrameric c-di-GMP mediates effective transcription factor dimerization to control *Streptomyces* development. Cell 158:1136–1147.

5. Jones SE, Pham CA, Zambri MP, McKillip J, Carlson EE, Elliot MA. 2019. *Streptomyces* volatile compounds influence exploration and microbial community dynamics by altering iron availability. mBio 10:e00171-19.

6. Shepherdson EMF, Netzker T, Stoyanov Y, Elliot MA. 2022. Exploratory growth in *Streptomyces venezuelae* involves a unique transcriptional program, enhanced oxidative stress response, and profound acceleration in response to glycerol. J Bacteriol 204:e00623-21.

7. Zhang X, Andres SN, Elliot MA. 2021. Interplay between nucleoid-associated proteins and transcription factors in controlling specialized metabolism in *Streptomyces*. mBio 12:e01077-21.

8. Shepherdson EMF, Elliot MA. 2022. Cryptic specialized metabolites drive *Streptomyces* exploration and provide a competitive advantage during growth with other microbes. Proc Natl Acad Sci 119:e2211052119.

9. Gust B, Challis GL, Fowler K, Kieser T, Chater KF. 2003. PCR-targeted *Streptomyces* gene replacement identifies a protein domain needed for biosynthesis of the sesquiterpene soil odor geosmin. Proc Natl Acad Sci U S A 100:1541–1546.

10. Gust B, Chandra G, Jakimowicz D, Yuqing T, Bruton CJ, Chater KF. 2004. λ Red-mediated genetic manipulation of antibiotic-producing *Streptomyces*. Adv Appl Microbiol 54:107–128.

11. Sherwood EJ, Hesketh AR, Bibb MJ. 2013. Cloning and analysis of the planosporicin lantibiotic biosynthetic gene cluster of *Planomonospora alba*. J Bacteriol 195:2309–21.

12. Gregory MA, Till R, Smith MCM. 2003. Integration site for *Streptomyces* phage ϕBT1 and development of site-specific integrating vectors. J Bacteriol 185:5320–5323.
